# An Enhanced Sampling Approach to the Induced Fit Docking Problem in Protein-Ligand Binding: the case of mono-ADP-ribosylation hydrolases inhibitors

**DOI:** 10.1101/2021.05.08.443251

**Authors:** Qianqian Zhao, Riccardo Capelli, Paolo Carloni, Bernhard Lüscher, Jinyu Li, Giulia Rossetti

## Abstract

A variety of enhanced sampling methods can predict free energy landscapes associated with protein/ligand binding events, characterizing in a precise way the intermolecular interactions involved. Unfortunately, these approaches are challenged by not uncommon induced fit mecchanisms. Here, we present a variant of the recently reported volume-based metadynamics (MetaD) method which describes ligand binding even when it affects protein structure. The validity of the approach is established by applying it to a substrate/enzyme complexes of pharmacological relevance: this is the mono-ADP-ribose (ADPr) in complex with mono-ADP-ribosylation hydrolases (MacroD1 and MacroD2), where induced-fit phenomena are known to be operative. The calculated binding free energies are consistent with experiments, with an absolute error less than 0.5 kcal/mol. Our simulations reveal that in all circumstances the active loops, delimiting the boundaries of the binding site, rearrange from an open to a closed conformation upon ligand binding. The calculations further provide, for the first time, the molecular basis of the experimentally observed affinity changes in ADPr binding on passing from MacroD1 to MacroD2 and all its mutants. Our study paves the way to investigate in a completely general manner ligand binding to proteins and receptors.

## Introduction

Structure-based drug design (SBDD) analyzes at the atomic level how ligands interact with their biological targets. It identifies new ligands by mapping biological activity data to distinct structural and energetic features of ligand-target complexes. *In silico* predictions of the poses and affinities of ligands for their biological targets constitute a key step of the approach [1, 2].

The predictions challenge the computational chemist when considerable structural adaptations of the ligand and the target are involved [3–5]. Indeed, induced-fit mechanisms play a role beyond the better ligand juxtapositioning in the binding site: slow targets’ conformational transitions induced by ligands during the binding/unbinding process govern the differences of the free energies of all states along the binding/unbinding pathway [6, 7]. These differences can determine ligand specificity even in structurally highly similar targets, with changes in affinity up to several hundred-fold [6]. Thus, the knowledge of those pathways can reveal induced-fit mechanisms and allow the calculation of other important pharmaceutical parameters like residence time [8–12].

Traditionally, in a drug design setting, medicinal and computational chemists often use molecular docking that has the enormous advantage of being computationally cheap [13]. It can screen poses and trends in affinities for 10^6^-10^7^ compound libraries in the order of days (with performances depending on docking software and rotational freedom of the compounds [14]). However, these approaches quantify target–ligand energies in terms of simple scoring functions [15–17], that generally neglect entropic contributions of the binding along with the solvent effects [18]. As a result, scoring functions can be used at most for qualitative comparisons. In addition, and most importantly for the present discussion, they cannot predict (un)binding pathways and oversimplify the binding process: usually only residues around the ligand are considered to be flexible, under the assumption - not justifiable with induced-fit - that neither the binding site nor the binding mode is changed significantly. Therefore, docking algorithm performance drops in the presence of induced fit even in predicting poses and affinity trends. These limitations can in theory be overcome by predicting (un)binding processes and affinities by means of molecular dynamics (MD) simulations. The main issue in this approach is given by the timescales in which these processes live (from the order of the microsecond and longer). Despite using an especially designed calculator (i.e., Anton) that allows the identification and characterization of the binding pose of a ligand inside a protein [19], a quantitative description is currently achievable only for fast binders [20], within the limitation of the force field.

For slower binders (i.e., with a residence time that exceeds the timescale reachable by specific purpose computers-namely milliseconds-) two classes of MD-based approaches can be used. One is based on free energy perturbation (FEP) approaches [21–23]. These predict relative differences in binding free energies of ligands for the same target, of the same ligands for a set of different targets or for mutants of the same target [24–28]. They correctly take into account entropic effects and they are used widely in drug design campaigns [29]. However, likewise molecular docking approaches, they cannot predict (un)binding pathways and become infeasible when significant induced-fit effects take place [19, 30].

These latter limitations may be overcome by potential mean force calculations. These predict the absolute binding/unbinding free energy as a function of reaction coordinate(s) or collective variable(s) (CVs) and provide binding/unbinding pathways. Among these methods, metadynamics (MetaD) has been shown to be particularly successful to predict unbinding pathways even in case where target flexibility plays a significant role [31–36]. However, in the presence of induced-fit mechanisms, the correct identification of the relevant CVs requires knowledge of the ligand binding/unbinding processes, in particular of the conformational rearrangement taking place upon binding. This is not trivial to know a priori when induced fit is involved. Volume-based MetaD might in principle solve this issue by using system-independent CVs allowing the ligand to explore all the accessible volume surrounding the target [37]. However, in practice the volume needed to exhaustively cover the relevant conformational space with induced fit (e.g. for including large induced conformational changes upon binding) is very large. As a result, recrossing events (required for the convergence of the free energy) are not very likely and the calculations are overall not very efficient.

Herein, we have improved volume-based MetaD by: (i) Combining the strengths of system-independent CVs with the benefits of reducing the ligand accessible conformational space to target smaller volumes relevant for the binding. This strategy was already implemented in other MetaD approaches to favor the recrossing events [38–41]. (ii) Restraining the part of the target that is not directly involved in the binding process. This reduces the chances to minimize the possibility of local unfolding caused by the presence of a localized potential. We show here that this new approach, called Localized Volume-based Metadynamics (LV-MetaD), does not lose the adaptability to conformational changes upon-binding, and it dramatically decreases the computational cost needed to obtain convergence. Furthermore, it permits the exploration of possible ligand binding pathways and it finally overcomes the challenges of ligand induced fit and slow conformational changes by predicting poses, affinities and binding mechanisms.

As a test case, we focus on the human MacroD1 and MacroD2 enzymes. These catalyze the hydrolysis of mono ADP-ribose (ADPr) from protein substrates [42] as well as they can efficiently reverse ADPr modification from the 5′ or 3′ terminal phosphates of DNA and RNA [43]. ADP-ribosylation is an important post-translational modification (PTM) that occurs in multiple key biological processes. In spite of the high pharmacological relevance of these targets [42], drug design campaigns have been unsatisfying [44, 45], mostly because they have not been able to identify selective inhibitors for the two enzymes. These would provide important tools to regulate intracellular signaling and regulatory processes, with important applications in neurodegeneration and cancer [46].

The X-ray structures of the two proteins with and without ADPr (*apo*- and *holo*-forms, respectively) show that the catalytic sites are deeply buried in the protein [47, 48]. Two active loops (loop1 and loop2 red in Figure 1a) delimit the binding site and function as switch loops to sequester the substrate and they provide sufficiently structural flexibility to accommodate diverse types of substrates [49]. Therefore, not only does ligand binding require substantial changes of the loops’ conformations to access in the inner cavity, but also it induces specific loops’ conformations upon binding that cannot be predicted *a priori* [50].

**Figure 1.**
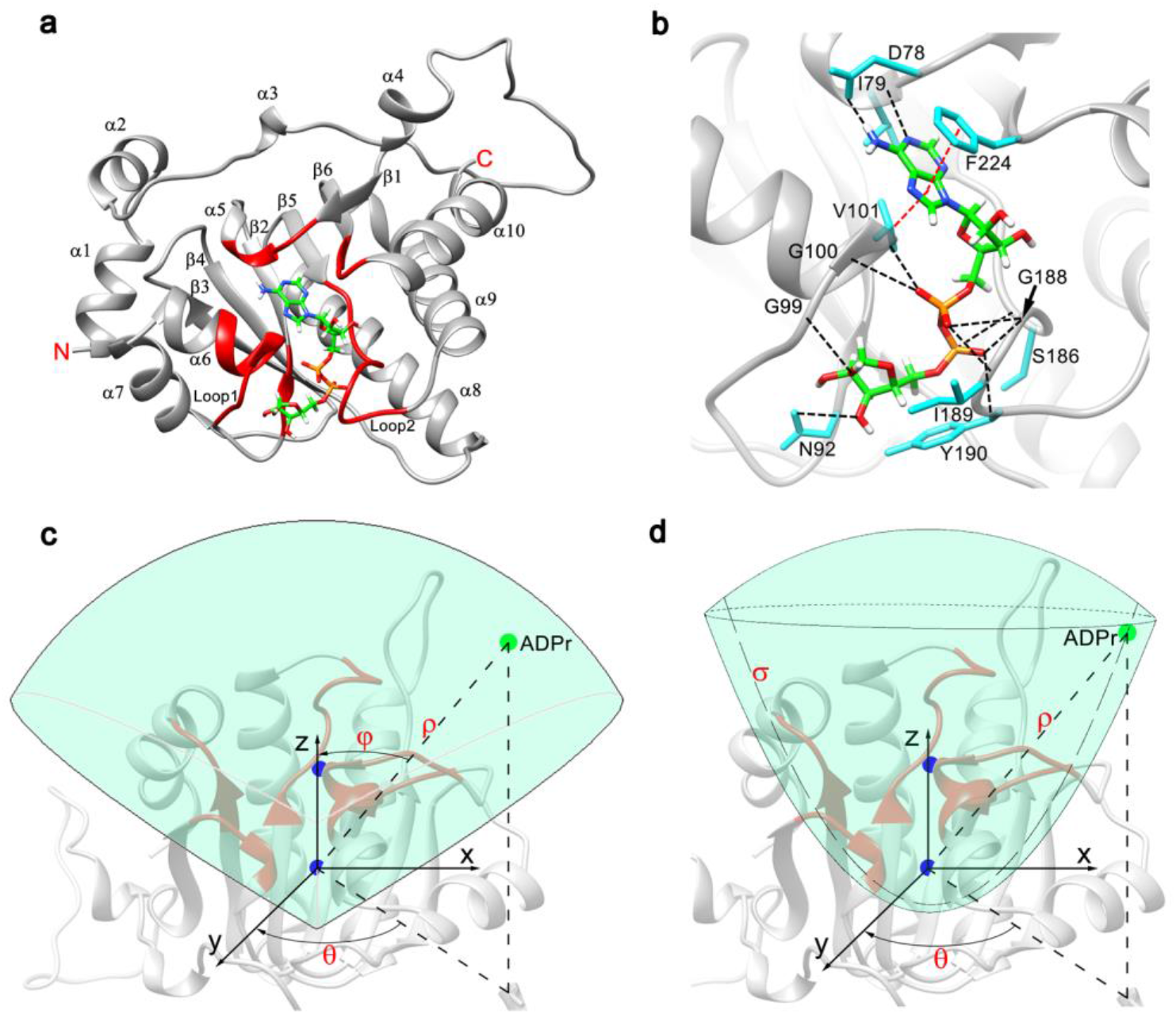
MacroD2 X-ray structure and the volume used in our Localized Volume-based Metadynamics simulations. (a) WT holo-MacroD2 X-ray structure (PDB code: 4IQY). The protein features a three-layered α-β-α sandwich-like conformation. The ligand ADPr is represented as sticks. The part of the protein involved in the binding pocket is highlighted in red. Loop1, 97-GGGGV-101, and loop2, 188-GIYG-191 are highlighted in red color. The backbone of the parts in gray undergoes a light constraint. (b) ADPr pose. Hydrogen bonds and hydrophobic interactions are depicted as black and red dashed lines, respectively. (c) ⅓ sphere-solid restraint volume. The ρ, ϑ, and φ spherical coordinates define ADPr position. (d) Parabolic-solid restraint potential. The ρ, ϑ and σ coordinates identify the position of the restraint potential. In both (c) and (d), the residues involved in the ligand binding site (in red) are completely contained in the restraining volume. The center of mass of the protein and binding site (shown in blue balls) are considered to define the orientation of the volume.

Inspection of the X-ray structures shows that in both enzymes, the ligand ADPr presents an L-shape conformation and the binding poses of the ligand are the same (Figure 1b and S1). The residues in the two binding sites are also the same on passing from MacroD1 to MacroD2, except that a phenylalanine (F272) and a valine (V271) in the former are replaced by a tyrosine (Y190) and an isoleucine (I189) in the latter, respectively (Figure S1). These two different residues in loop2 cause a change in affinity toward ADPr from −9.5 kcal/mol to −8.4 kcal/mol on passing from MacroD2 to MacroD1, respectively. Further mutations of these same two key residues in MacroD2, Y190N and I189R, substantially increase the affinity for ADPr up to −10.1 kcal/mol and −10.3 kcal/mol, respectively [50]. This experimental evidence suggests a key role of the loops’ residues in substrate specificity. However, the molecular mechanisms of such changes in affinity are not known.

Our LV-MetaD scheme turned out to reproduce (i) the ADPr binding poses of the known complexes in the wild-type (WT) systems and it predicts the unknown ones for Y190N and I189R MacroD2 mutants. (ii) the experimentally measured protein-ADPr binding free energy of the two WT enzymes, along with those of the Y190N and I189R MacroD2 mutants, providing a rationale for the increased affinity of the latter. This information may help understand the druggability and substrate specificity of such classes of proteins. Furthermore, our calculations elucidate the molecular interactions induced upon binding at each stage of the binding/unbinding process, paving the way to the design of highly selective inhibitors. Finally, our calculations provide the molecular basis, for the first time, of the impact of all the mutagenesis experiments on the measured binding affinities so far conducted. Overall, our novel LV-MetaD protocol emerges as a powerful tool to efficiently investigate the induced-fit molecular recognition in ligand/receptor events.

## Results and Discussion

LV-MetaD calculates the ligand substrate (ADPr) binding free energies as a function of apt collective variables. These are the 3D positions of the ligand relative to the binding pocket. The calculations allow to predict substrate ADPr poses and binding pathways in WT MacroD1 and WT, Y190N and I189R MacroD2. An efficient sampling is achieved by: (i) Limiting the exploration of the ligand within a well-defined volume around the target binding pocket, thus avoiding that most of the sampling time for the ligand is spent in the solvent, a portion of the space not relevant for binding and unbinding. This is achieved by adding a repulsive potential at the boundaries (see Section below). (ii) Adding a constraint on the backbone in parts of the protein not directly involved in the binding/unbinding process (red part in Figure 1c and 1d). This potential is added because the ligand can be forced to explore volume portions occupied by the host, unfolding the protein.

### Choice of the volume

For WT MacroD2, we set up 2 different volume shapes to limit the sampling of the ligand. These are either a ⅓ sphere-solid shape (Figure 1c) or a parabolic-solid (Figure 1d and Figure S2) shape. The exploration of the conformational space is satisfactory as several recrossing events occur in both cases (Figure S3). The binding free energies, calculated as specified in the Methods section, turns out to agree well with experimental measurement (Tables S1). This confirms the flexibility of the method irrespective of the choice of the volume shape; indeed, the latter can be adapted to maximize sampling efficiency depending on the specific case.

Here, since we observed a slightly better agreement with the experimental data (Table S1) for parabolic-solid volume in MacroD2, we decided to implement this shape for all the remaining systems. The latter allowed us to compute binding affinities in extremely good agreement with the experimental ones (Table 1).

**Table 1.** Absolute free energy from parabolic-solid LV-MetaD simulation (ΔG_standard) and experiment [49–51] (ΔG _exp).

| Systems | $\Delta G_{\text{metad}}$<br>(kcal/mol) | $\Delta G_{\text{correction}}$<br>(kcal/mol) | $\Delta G_{\text{standard}}^a$<br>(kcal/mol) | $\Delta G_{\text{exp}}^b$<br>(kcal/mol) |
| --- | --- | --- | --- | --- |
| MacroD2 | -7.8±0.3 | 1.6 | **-9.4±0.3** | **-9.5** |
| MacroD1 | -6.6±0.3 | 1.6 | **-8.2±0.3** | **-8.4** |
| MacroD2 I189R mutant | -8.1±0.3 | 1.6 | **-9.7±0.3** | **-10.3** |
| MacroD2 Y190N mutant | -8.4±0.3 | 1.6 | **-10.0±0.3** | **-10.1** |

$\Delta G_{\text{standard}}^a$ is calculated at $T = 298$ K and the ionic strength of 100 mM. $\Delta G_{\text{exp}}^b$ of WT MacroD2 and its mutant is determined at 298 K, while $\Delta G_{\text{exp}}^b$ of WT MacroD1 is detected at 303 K. The ionic strength is 100 mM and the method of binding assays is isothermal titration calorimetry.

All the systems were initially equilibrated by 300 ns of MD, followed by ~0.8 μs-long LV-MetaD simulations. In the next Sections, we discuss the impact of induced fit on the binding poses and the (un)binding processes for each system.

#### WT MacroD2

The calculations on the MacroD2 were based on the *holo* structure of MacroD2 in complex with ADPr [52]. The free energy shows the presence of four basins (Figure 2). The lowest free energy minima (basins **1** and **2**) differ by only 1 kT (thus they are both populated at room temperature) and represent the bound state. The structure associated with basin **2** reproduces the pose in *holo-* MacroD2 X-ray structure (Figure 3a and 3b) [50]: the ADPr *adenine* ring is deeply buried in the highly conserved binding pocket [50] and forms bifurcated H-bonds with the backbone and sidechain of D78, respectively. The ADPr *adenosine ribose* forms H-bonds with the backbones of C222 and C184. The negatively charged ADPr *pyrophosphate* moiety located in the cleft of active loop1 and loop2 forms H-bond networks with backbone atoms of I185, S186, T187, G188, I189, and bifurcated H-bonds with the backbone and sidechain of Y190. The ADPr *distal ribose* faces the entrance of the binding cavity. It forms H-bonds with backbone of A90 and sidechain of N92 in loop1.

**Figure 2.**
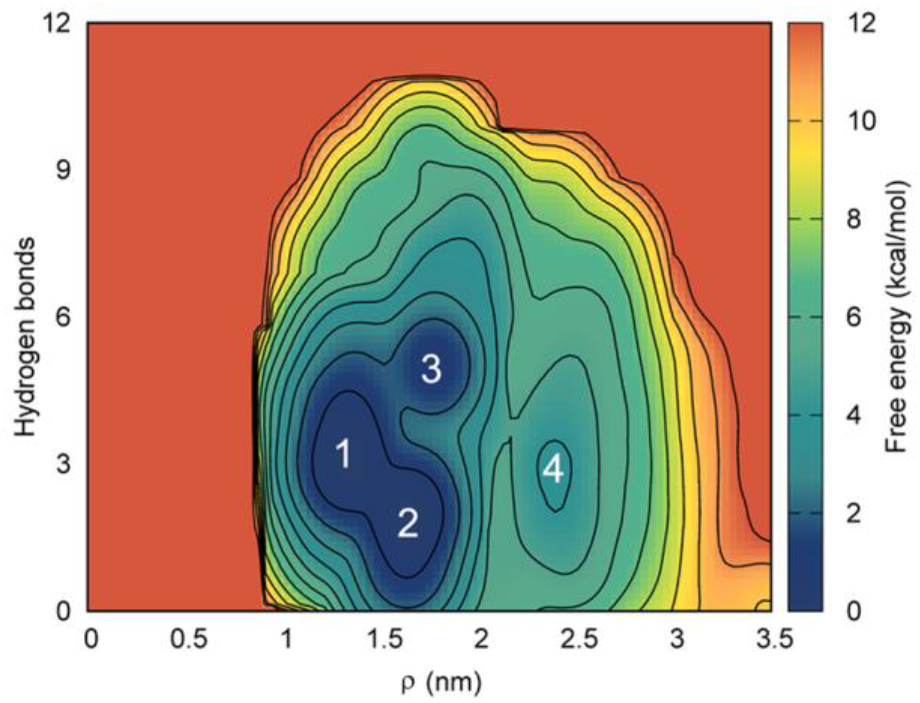
Free energy surface associated with MacroD2/ADPr binding as a function of ρ (the distance between the center of mass of the target and the center of mass of the ligand and the number of hydrogen bonds between the residues in the binding pocket and the ligand (n_H). In LV-MetaD, free energy as calculated in LV-MetaD as a function of the spherical or parabolic coordinates of the ligand, cannot be interpreted straightforwardly. To overcome this issue, the figure shows the free energy surface reweighted along variables that can be more meaningful for the identification of relevant minima, such as ρ and the number of hydrogen bonds.

**Figure 3.**
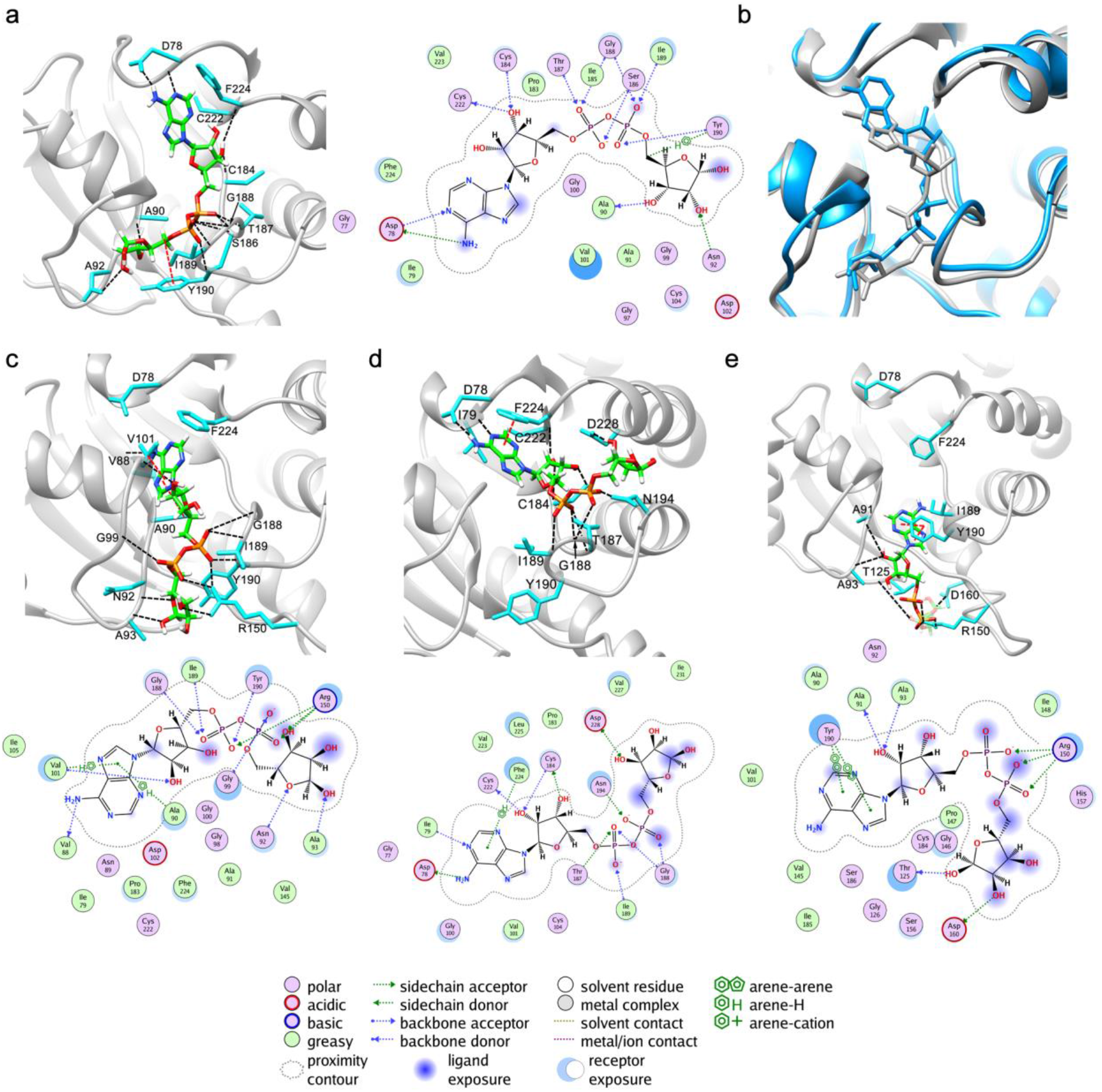
LV-MetaD simulation of ADPr binding to WT MacroD2. (a) Representative conformations of basin **2** (left) and its ligand-protein interaction diagram (right). Bulk water is not shown for clarity Hydrogen bonds and hydrophobic interactions are depicted as black and red dashed lines, respectively. (b) Superposition of holo-MacroD2 X-ray structure (in grey) with the binding pose of basin **2** (in blue). (c), (d), (e) Same as (a) for basin **1, 3, 4.** The calculations use the parabolic-solid scheme of Figure 1.

In basin **1** (Figure 3c), the *adenine* ring of ADPr no longer Interacts with D78 and F224 due to slight variation of ADPr pose. These interactions are substituted by the formation of hydrophobic contacts with residues V101 and A90 and H-bonds with V88 backbone (Figure 3c). ADPr *adenosine ribose* no longer forms hydrogen bonds interactions with the backbone of C222 and C184 but with the backbone of V101. The *pyrophosphate* moiety form hydrogen bonds with the sidechain of R150 and backbone atoms of residues in the loop2 (i.e. G188, I189, and Y190) as well as with G99 (Figure 3c), which is similar to what is observed in basin **2**. The *distal ribose* faces the solvent and forms hydrogen bonds with the backbones of N92 and A93, respectively.

In basin **3** (Figure 3d) the ADPr is partially out from the cleft between the two loops and exposed to the solvent. The *adenine* ring maintains hydrogen bonds with the sidechain of D78 and backbone of I79 and a π-stacking with F224, and the *adenosine ribose* forms the H-bonds with the C222 and C184 backbones and with C184 sidechain (Figure 3d). But the *pyrophosphate* and *distal ribose* moiety rotate above loop2 and are partly sandwiched between helix9 and helix10. The *pyrophosphate* moiety is stabilized by the H-bonds with T187, G188, I189 and N194, while the *distal ribose* loses all the interactions with loop1 and loop2. In basin **4**, the ADPr is located at the entrance of the binding cavity (Figure 3e). The *adenine* ring is located in the cleft formed by the loop1 and loop2 and forms π-stacking interactions with Y190 (Figure 3e), and the *adenosine ribose* forms H-bonds with backbones of A91 and A93. The *pyrophosphate* moiety and distal ribose is completely exposed to the solvent, the former forms H-bonds with R150 sidechain, while the *later* forms H-bond interactions with T125, and D160, backbone and sidechain, respectively.

From this free energy landscape, we can gain qualitative insight on the induced-fit ligand (un)binding mechanism: the ADPr *adenine* ring approaches the binding pocket by interacting with loop1, then it passes through the cleft between loop1 and loop2 and subsequently reaches the inner binding site in the binding pose (Figure S4b). During the unbinding process, the *pyrophosphate moiety* and *distal ribose* of ADPr escape from the groove between loop1 and loop2 and move towards the cleft between helix 9 and helix 10 and then lose all the interactions with the binding pocket (Figure S4b). In the bound state (basin **2**), the loops are as close as 10.8 Å (centroid distance). The transition from the bound to the unbound state, however, triggers a closure and an opening of the loop1-loop2 interface (Figure S4c and S4d), with their centroid distances ranging from a minimum value of ~10.0 Å to a maximum of ~14.5 Å (Table S2 and Figure S4c), indicating the presence of ADPr dependent induced-fit mechanisms. Such movements of the loops cannot be appreciated in the X-ray structure where the loop conformations in *apo* and *holo* states are very similar, with centroid distances of 12.9 Å and 11.6 Å, respectively (Table S2) [52]. Therefore, our simulations reveal key ligand-dependent events involving loop1 and loop2 impacting on the binding pathway, and in turn on the binding affinity (see follow-up paragraphs).

#### WT MacroD1

The calculations on WT MacroD1 were based on the *apo* X-ray structure instead of the *holo* structure (Figure 4a) to check the robustness of the method in reproducing induced-fit effects of the binding process even starting from a conformation where the ligand is not at its energy minimum. As already discussed above, they turned out to reproduce the free energy of binding (Table 1).

**Figure 4.**
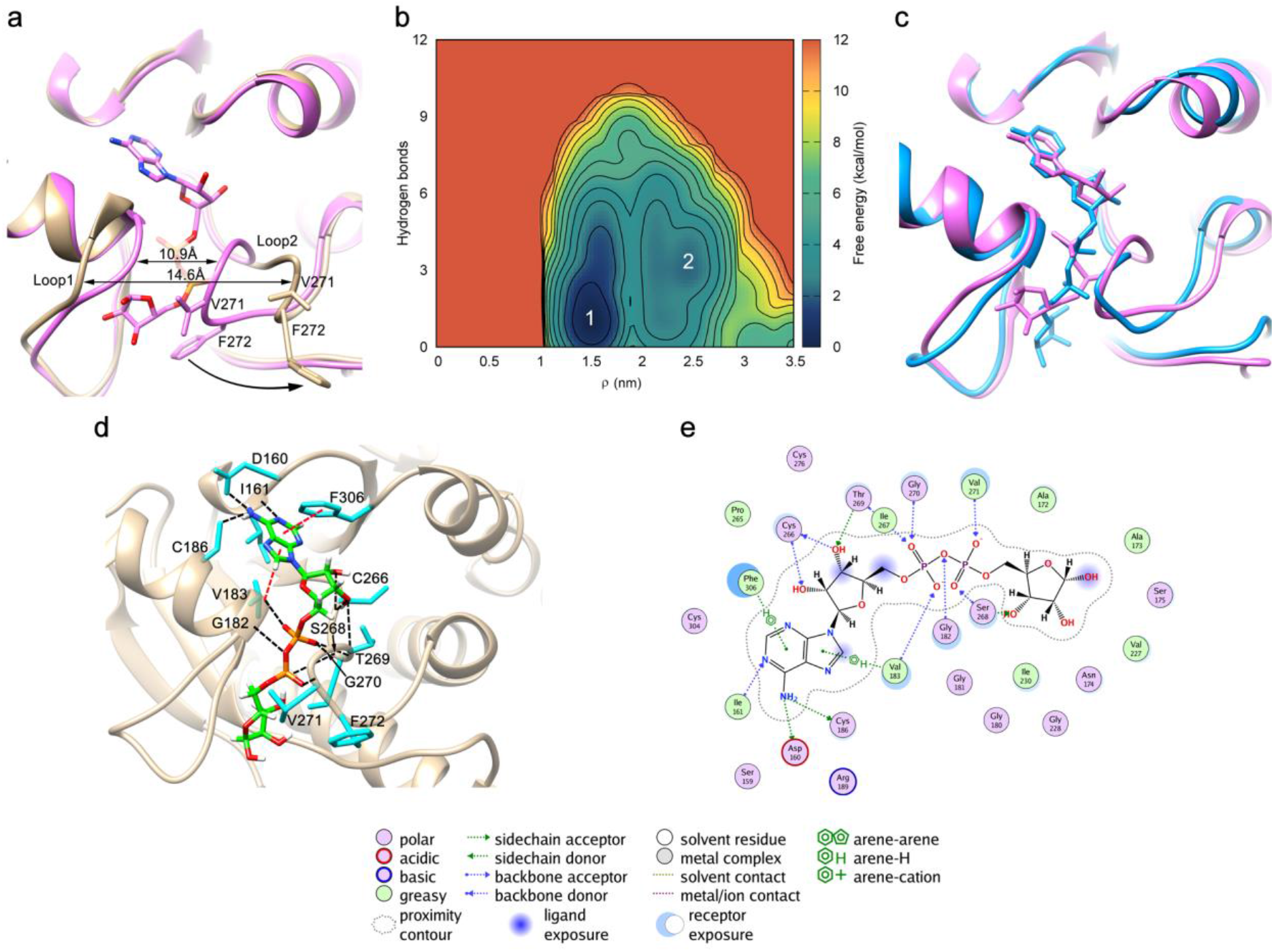
LV-MetaD simulation of ADPr binding to apo-MacroD1. (a) Superposition of holo-MacroD1 (PDB code: 6LH4, in purple) and apo-MacroD1 (PDB code: 2X47, in yellow) X-ray structures. The distance between the center of mass of loop1 and loop2 are highlighted. They consist of 179-GGGGV-183 and 270-GVFG-273 residues, respectively. (b) Free energy surface as a function of ρ and n_H. (c) Superposition of the holo-MacroD1 X-ray structure (in purple) with the calculated MacroD1_ADPr binding pose of free energy basin **1** (in blue) highlighted on the free energy landscape. (d) Representative conformation of free energy basin **1**. Same color-code and representation mode of Figure 3 is used; (e) Ligand-protein interaction diagram for representative structure of free energy basin **1**.

The free energy landscape features only two basins, **1** and **2** (Figure 4b). Remarkably, basin **1** corresponds to the pose observed in the *holo* MacroD1 X-ray structure (Figure 4c). It features a close conformation as in the *holo form*, where the centroid distances of loop1 and loop2 are ~10.5 Å and ~10.9 Å, respectively (Figure 4a and Table S2) [49], to be compared with the distance in the *apo* form where loop1 and 2 are in an open conformation with a centroid distance of ~14.6 Å. This is a remarkable result, as the simulation started from the *apo* form with the ligand docked a posteriori, instead of the *holo* form. In basin **1**, the *adenine* ring of ADPr binds to the inner binding pocket establishing π-stacking interaction with F306 and maintains a similar hydrogen bonding network as in the *holo*-MacroD1 structure (Figure 4d and S5c). The *adenosine ribose* forms H-bonds with the backbone of C266 and sidechain of T269. The *pyrophosphate* moiety is stabilized by hydrogen bonds formed with backbones of the residues in the two active loops, i.e. G182, V183, S268, T269, G270 and V271 (Figure 4e). The *distal ribose* is slightly further away from loop1 and it does not form direct interactions with the two loops. Indeed, it is slightly twisted with respect to the *holo-* MacroD1 and MacroD2 X-ray structure (Figure 4c). Basin **2** is ~ 4 kcal/mol higher in free energy with respect to basin **1** (Figure 4b). Here, the *adenine* ring of ADPr is stacked between loop1 and loop2 (Figure S6a). The other parts of the ligand are completely exposed to the solvent above loop1, forming hydrogen bonds with the backbone amino groups of G182, G185 and C186, and with the sidechain guanidino group of R189 (Figure S6b).

Also in this case, we attempt to gain insight on the (un)binding process by investigating the free energy landscape. The loop distance change increases relative to the WT MacroD2: the centroid distance ranges between 10.3 Å and 16.8 Å (Table S2). The *adenine* ring of ADPr initially approaches the binding site by interacting with helix5 or helix10 and then the other moieties of the ligand directly lie in the groove between loop1 and loop2. During the unbinding process, the ligand moves its *distal ribose* and *pyrophosphate* moiety towards the solvent and then drags the *adenine* moiety directly from the cleft between the loop1 and loop2 (Figure S5b). This is relevant if we consider that the two residues that discriminate between MacroD2 to MacroD1 (F272 and V271 in MacroD1 are Y190 and I189 in MacroD2) are located in loop2: these difference in the amino acid sequence indeed affect the steric hindrance of the entrance of the binding pocket (V271 in MacroD1 is smaller than I189 in MacroD2), as well as the number of protein-ligand interactions (e.g. the H-bonds between *pyrophosphate* moiety and F272 are not present in MacroD1), and in turn the ADPr-induced loops’ configurations, possibly providing a rationale also for the lower number of intermediate states. Thus, the two residues which are modified on passing from WT MacroD1 to WT MacroD2 cause a loss of interactions between ADPr and loop2, leaving loop1 as the sole player in ligand binding, in contrast to WT MacroD2. Again, these aspects were not identifiable by just the comparison of the *apo* and *holo* crystal structure of MacroD1.

#### Y190N and I189R MacroD2

No structural information is available for these systems, therefore, they were modeled from WT *holo* MacroD2 (see Methods).

#### Y190N MacroD2

The free-energy profile shows four basins as in the WT protein (Figure 5a). Basin **1** is the lowest free energy minimum. The ligand pose is similar to those of WT MacroD2/1 (Figure 5b). The interactions established by the *adenine* ring and the *adenosine ribose* with the protein are similar to the ones established in MacroD2 (basin **2**). The *distal ribose* and *pyrophosphate* moiety form extensive hydrogen bond networks with residues in the vicinity of loop1 and loop2, i.e. backbones of G100, V101, S186, T187, G188 and sidechain of D102 (Figure 5c and 5d). However, the *pyrophosphate* moiety no longer interacts directly with I189 and N190 as in WT MacroD2. Indeed, we observed that the active loop2 in Y190N MacroD2 is in an open conformation, different from the close one observed in the WT MacroD2 bound state (Figure 5b). In basin **2**, the ligand locates at the entrance of the binding site appearing as in a pre-binding state (Figure S7a), while in basin **3**, ADPr hangs over the gap between loop1 and loop2 (Figure S7b). In basin **4**, ADPr completely escapes from the binding pocket (Figure S7c). Y190N MacroD2 displays the same number of basins found in WT MacroD2. However, interestingly, during the binding/unbinding process the centroid distances between the two loops sample a broader range of values (from 10.2 Å to 22.5 Å, Table S2) with respect to the WT MacroD2. Notably, the two loops are all in an open conformation in basins **1-4**. Also, ADPr undergoes the same binding process as WT MacroD2 but a different unbinding process. Namely, ADPr escapes from the binding site along the cleft between helix5 and helix6 (Figure S8b). These differences on passing from WT MacroD2 to the Y190N mutant might be caused by the smaller sidechain in the 190 position, which from one side increases loop2 flexibility and from the other side decreases the steric hindrance of the cleft between loop1 and loop2. The observed effects could provide a rationale for the increased binding affinity.

**Figure 5.**
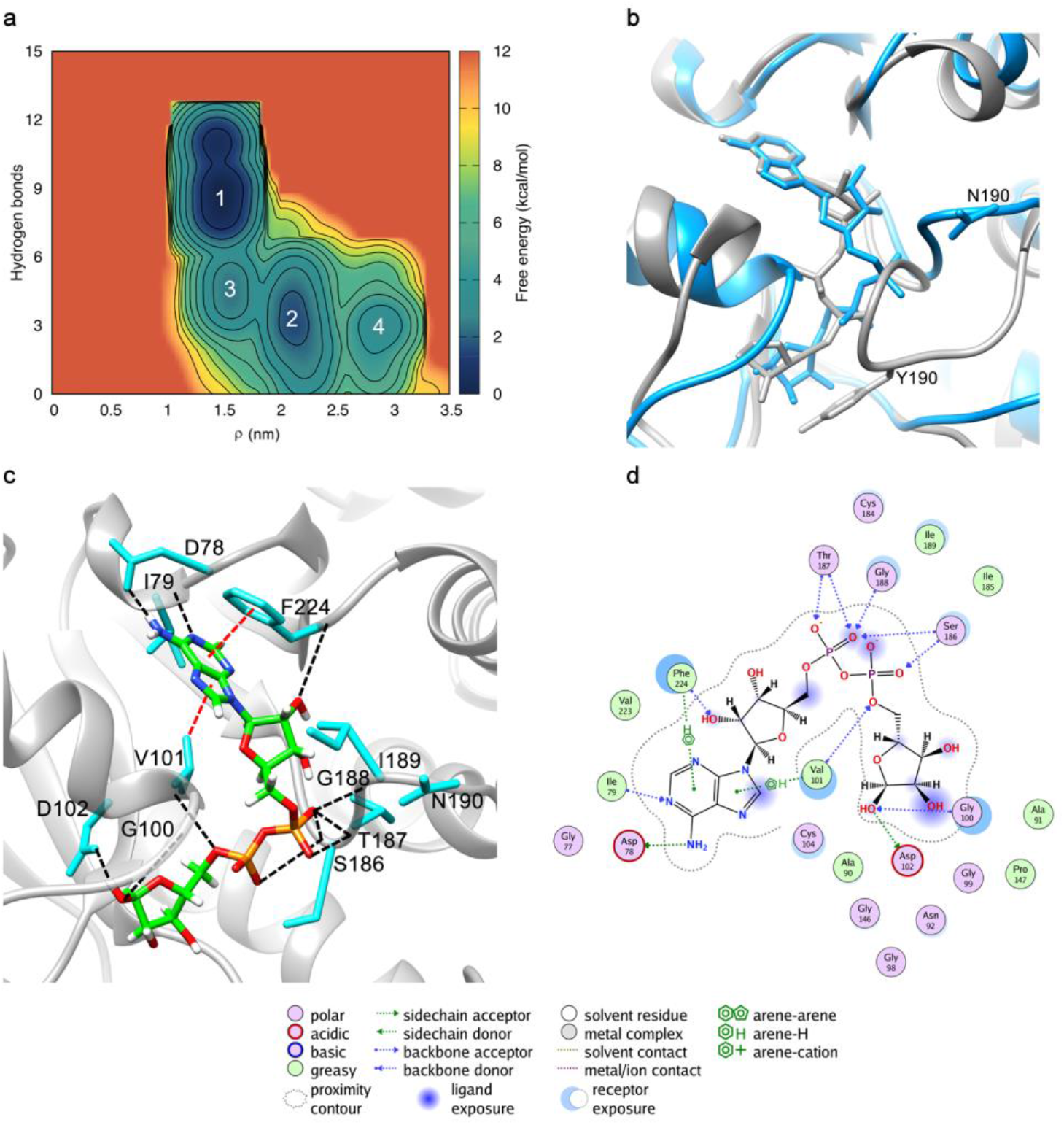
LV-MetaD simulation of ADPr binding to the Y190N MacroD2. (a) Free energy surface as a function of ρ and n_H. ρ is here the distance of the center of mass between Y190N MacroD2 and ADPr. (b) Superposition of WT holo-MacroD2 X-ray structure (in gray) with the binding pose of free energy basin **1** (in blue) highlighted on the free energy landscape. (c) The representative conformation of free energy basin **1** in the binding pocket. Same color-code and representation mode of Figure 3 is used. (d) Ligand-protein interaction diagram for representative structure of free energy basin **1**.

#### I189R MacroD2

Only three basins are observed (Figure 6a), in contrast to four of the WT MacroD2 and the Y190N mutant. This might be due to the fact that R189 attracts the ligand and locks its orientation inside the binding pocket. Therefore, the conformational freedom of the ligand in the pocket is reduced, possibly lowering the number of intermediate states. The binding pose of ADPr in basin **1** reproduces the conformation observed in WT MacroD2 and involves almost the same interactions (Figure 6b). The main difference is that the negatively charged *pyrophosphate* moiety is stabilized by a salt bridge interaction with R189 (Figure 6c and 6d). This new interaction might contribute to the observed higher ADPr binding affinity. Basin **2** corresponds to a pre-dissociated transition state where the ligand attempts to escape from the binding groove into the solvent by moving its distal ribose across loop1. During the motion, the salt bridge interaction with R189 is however maintained (Figure S9a). The binding pose of basin **3** occurs when the ligand leaves basin **2** to reach a more solvent exposed state (Figure S9b). During the binding/unbinding process, the centroid distance between loop1 and loop2 varies from a maximum distance of ~17.5 Å to a minimum distance of ~9.8 Å, having 10.6 Å in the bound state (basin **1**), comparable to the value observed in the bound states of ADPr in the WT MacroD2.

**Figure 6.**
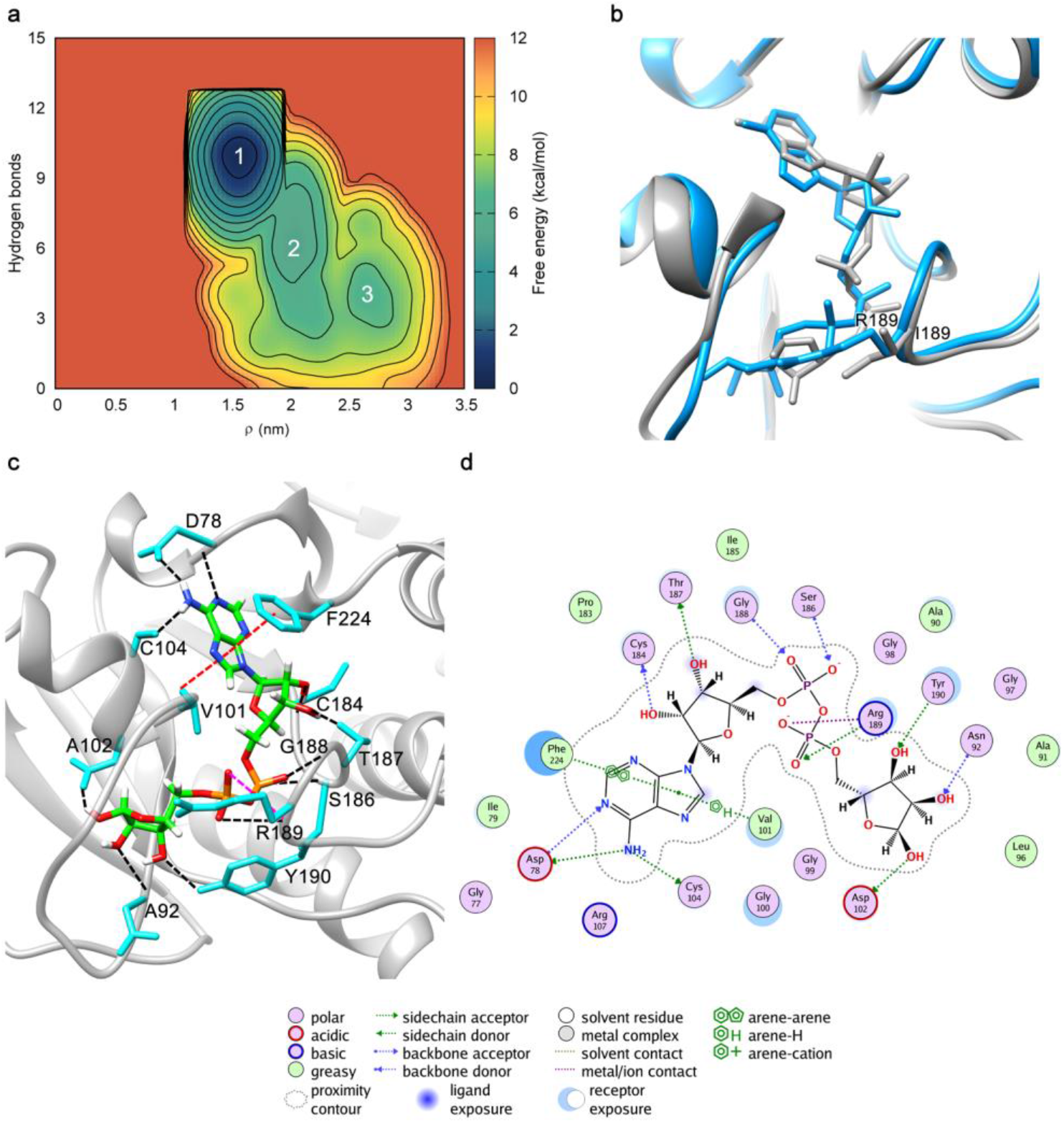
LV-MetaD simulation of ADPr binding to the I189R MacroD2. (a) Free energy surface as a function of ρ and n_H. ρ is the distance of the center of mass between I189R MacroD2 mutant and ADPr. (b) Superposition of WT MacroD2 crystal structure in complex with ADPr (in grey) with the binding pose of free energy basin **1** (in blue) highlighted on the free energy landscape. (c) The representative conformation of basin **1** in the binding pocket. Same color-code and representation mode of Figure 3 is used. (d) Ligand-protein interaction diagram for representative structure of free energy basin **1**.

#### Molecular basis for the change in affinity of MacroD2 mutants

Based on the affinity of ADPr, the Macro2 mutants have been classified in three groups [50]: **(i)** variants featuring low affinity (below 20% of that of the WT); **(ii)** variants featuring intermediate ADPr affinity (about 40-60 % of that of the WT); **(iii)** variants featuring WT-like activity and affinity for ADPr or even higher than the WT, i.e. I189R and Y190N, investigated above. On the basis of our calculations, we attempt here to provide a rationale for the changes in affinity of all the mutants by visual inspection of the two free energy minima of WT MacroD2/ADPr (basins **1** and/or **2** in Figure 3), and of the X-ray structures (Figure S1b).

G188E, G100E and G100E/I189R MacroD2 belong to **(i).** In basin **2** and in the X-ray, G188 backbone establishes bifurcated H-bonds with the negatively charged *pyrophosphate* moiety of ADPr; G100 backbone establishes a single H-bond with the *pyrophosphate* moiety in the X-ray, while in basins **1** and **2**, it is in close proximity (Figure 3a). Therefore, the G100E and G188E mutations are expected to introduce a bulky negatively charged group near to *pyrophosphate*, leading to the disruption of the hydrogen bonding network between the *pyrophosphate* and the two loops. This may lead to rearrangement of how the ligand is positioned in the catalytic site or it might even prevent binding completely, possibly explaining the dramatic loss of affinity. The affinity of G100E/I189R MacroD2 is larger than that of G188E and G100E MacroD2 but still lower than that of variants in group **(ii)**. The I189R mutation introduces a positively charged group close to the negatively charged *pyrophosphate*. Thus, we speculate here that the I189R mutation stabilizes, at least in part, loop2/ligand interactions, leading to a less dramatic change of affinity relative to the WT, as observed experimentally.

D78A, G100S and N92A/D102A MacroD2 belong to **(ii)**. In basin **2**, D78 sidechain and backbone establish bifurcated H-bonds with ADPr’s amino group N6 and N1 atoms (Figure 3a), respectively. The H-bond with N6 is observed also in the X-ray structure (Figure S1). The N92 sidechain forms a H-bond with an OH group of the *distal ribose* in both basin **2** and in the X-ray (Figure 3a and Figure S1). These interactions are however not observed in basin **1** of the same system (where the binding pose of the ligand is still preserved), suggesting that these *aminoacids* are not key for stabilizing ligand binding. D102 in our simulations and in the X-ray is in close proximity to the *distal ribose* and the *pyrophosphate* moiety, respectively, but it does not appear to form key interactions with the ligand. Therefore, it is not possible to understand whether the D102A mutant might have an effect if uncoupled from the N92A. G100, as said above, only establishes a single H-bond with the *pyrophosphate* moiety in the X-ray that is not observed in basins **1** and **2**. Indeed, G100S has only a mild effect on the binding affinity. The smaller impact of G100S with respect to G100E described above can be rationalized by the smaller steric hindrance and missing net charge of serine with respect to glutamate.

I189R Y190N, F224A and G99E MacroD2 belong to **(iii).** The rationale for the increased affinity of I189R and Y190N has been given above. F224A and G99E MacroD2 feature affinities similar to that of the WT. Notably, while in the X-ray these residues feature a π-stacking with the *adenine* ring and H-bond with the distal *ribose*, respectively. These interactions are not observed in basin **2** and **1** of our simulations. G99 is indeed solvent-exposed in our simulations and therefore its substitution for E is not expected to affect ligand binding, as confirmed by the experiments (Figure 3a). F224 is only in close proximity to the ligand, therefore its substitution with A is not expected to play a key role for the affinity, consistently with experimental evidence.

Overall, our simulations can provide a rationale for the different affinity observed in mutagenesis experiments, not fully emerging just from the visual inspection of the X-ray structure.

## Conclusions

We have presented a new method, called Localized Volume-based MetaD scheme (LV-MetaD), to predict poses, affinities and (un)binding mechanisms of small ligands targeting proteins under *any* circumstances, even in case induced-fit effects are significant. The method exploits the advantages of system-independent CVs with a restricted conformational space sampling. To the best of our knowledge, this is the first time that such a general scheme has been developed. LV-MetaD has been applied to substrate (ADPr) binding to macrodomain enzymes, WT MacroD1 and WT MacroD2 and their binding to ADPr, for which we know that induced fit effects are present: two highly flexible loops regulate the access to the active site and rearrange in a specific manner depending on the ligand [49]. Simulations have been extended also to the Y190N MacroD2 and I189R MacroD2 mutants, which are the only mutants for which the affinity increases.

The calculated absolute binding free energy differences of the ligand binding to the WT proteins turn out to compare well with experiments. The scheme reproduces also the increased affinity of the mutants with respect to the WT MacroD2, although the difference in affinity between Y190N MacroD2 and I189R MacroD2 is beyond the precision of the method. Taken together, these results establish the accuracy of the calculations.

The calculations further predict the ligand poses of ADPr in MacroD2 mutants, without prior knowledge of it and depict the differences in the binding/unbinding pathways with respect to the WT MacroD2. In addition, they provide a rationale for the impact of the other mutations for their consequences of affinity. Overall, our simulations confirm that the conformational rearrangements of the binding pocket are induced by ADPr binding and they demonstrate the capability of LV-MetaD protocol to exactly calculate the binding free energy, predict the binding conformation and characterize the induced-fit binding mechanisms. Moreover, no prior knowledge of the binding pose is required.

The scheme is entirely general and it could be applied to investigate ligand binding to other enzymes and receptors for which induced fit is important. The free energy is described by a set of collective variables that can be set up according to the shape of the binding pocket. The application of this scheme with new and efficient form of CV-based enhanced sampling protocols, such as the on-the-fly probability (OPES) [53]), combined with the new computational capabilities given by the exascale computing project, can finally pave the way for a totally general *in silico* drug discovery approach based on molecular simulations.

## Supporting information

supplementary meterials

## Acknowledgment

The authors acknowledge Prof. Michele Parrinello for inspiring this work with his research and valuable suggestions, as well as for carefully reading the manuscript. The authors thank Somayeh Asgharpour and America Luz Chi Uluac for fruitful discussions. This work is financially supported by the 2018-HGF-OCPC Program (China and Germany postdoctoral Exchange Program 2018) to GR, JL and BL. GR and PC acknowledge the (2020-2022) Helmholtz European Partnering program. J.L. acknowledged the financial support from the Natural Science Foundation of Fujian Province (2019J06007).

## Data Availability

All the data and PLUMED input files required to reproduce the results reported in this paper are available on PLUMED-NEST (www.plumed-nest.org), the public repository of the PLUMED consortium [54] as plumID: 21.018.

## Methods

### System preparation

The crystal structure of human mono-ADPr hydrolase macrodomains have been reported in the literature: the first is MacroD2 in complex with ADPr (PDB access code 4IQY), resolved at 1.5 Å atomic resolution [50]. The crystal structure has two protein molecules, by aligning them, the RMSD is less than 0.1 Å, revealing it has two nearly identical structures. Hence, in order to reduce the computational cost, we performed all the simulations using the monomer molecule. The second is MacroD1 monomer in the *apo* state at 1.7 Å resolution (PDB access code 2×47) [48] and MacroD1_ADPr complex at 2.0 Å resolution (PDB access code 6LH4) [49]. The two monomers involve a general macrodomain fold as three-layered α-β-α sandwich structure with a central six-stranded β sheet and are evolutionarily conserved protein with an almost identical binding site. In order to model a possible bound-conformation of ADPr, the initial structure of the MacroD1 system was built by docking ADPr to the apo-MacroD1 instead of starting from the holo-MacroD1. Additional two mutants of MacroD2 were obtained by replacing I189 with arginine and Y190 with asparagine respectively using UCSF Chimera [55]. The missing loops in these two protein crystallographic structures were reconstructed by using the protein preparation wizard of Schrödinger module [56]. Due to the lack of the structure of MacroD2 mutant bound to ADPr, the structure of WT MacroD2 complex was employed as a reference model for the MacroD2 mutant binding mode.

Each protein-ADPr complex system was performed with AMBER ff99SB-ILDN force field for the protein and ions [57], and was solvated with TIP3P explicit water molecules in a periodic cubic box, in which the protein surface are 20 Å far away from the periodic box edge[58]. Additional appropriate number of sodium and chloride counterions were added to neutralize the charge of the whole system and to achieve a physiological salt concentration of ~100 mM which was consistent with experimental ionic force used to determine the binding constant [50, 51]. Topology for the ligand was constructed using the general AMBER force field (GAFF) [59] and its partial atomic charges were parameterized with AM1-BCC semiempirical method [60].

### MD simulations

All of the simulations were performed using GROMACS 2019.2 [61] patched with the PLUMED 2.6 [54, 62]. Long-range electrostatic interactions were calculated with the particle mesh Ewald (PME) scheme using a grid spacing of 1.2 Å and the distance cutoff for the short-range electrostatic interactions and van der Waals was set to 12 Å [63]. All the bonds including the hydrogen bonds were restrained with the LINCS algorithm [64]. The velocity rescale thermostat [65] with solvent and solute coupled to separate heat baths was employed for the temperature regulation and the pressure was controlled by the Parrinello-Rahman barostat [66]. After an energy minimization of the solvent with the steepest descent algorithm, the temperature of each system was gradually increased from 0 to 298 K in 1 ns of MD. Keeping the position restraints for protein and ligand, the system was equilibrated at 298 K for 10 ns in the NVT ensemble; after this step we released the ligand, and the system was equilibrated by 10 ns in the NPT ensemble. Finally, a long equilibration of 300 ns without any restraint was performed before the production MetaD simulation.

### Metadynamics simulations

Here we developed the Localized Volume-based Metadynamics scheme on the base of funnel-shaped MetaD and spherical volume-based MetaD. We chose two different restraining volume shapes. The first restraining applied was ⅓ sphere-solid. As in the original volume-based protocol, we biased the spherical coordinate system: CV1-*ρ*, the distance between the center of mass of the ligand and protein, CV2-ϕ, the polar angle measured from the z-axis, and CV3-θ, the azimuthal angle of its orthogonal projection on the x-y plane [37, 67] (Figure 1c). To find an appropriate size of the restraining potential (i.e., that contains a reasonably solvated set of conformation for the ligand), we first performed a calculation to estimate the distance between protein and ligand to avoid any host-guest interactions. After this estimate, the value of *ρ* was restricted to 35 Å and the φ was limited to pi/3. The second restraining potential was shaped as a rotational parabolic-solid. Like in the first case, we biased three CVs, *ρ*, θ, and σ, where *ρ* and θ was the same spherical coordinate as above, and σ was a series of confocal parabolas centered on the center of mass of protein heavy atoms (Figure 1d). The value of σ (σ = 1) depends on the restraining space which should totally include the protein pocket and allow the ligand flexible exploration of the binding/unbinding orientation.

As previously said, a repulsive potential was applied at the edge of the designed volume, impeding the ligand from visiting less relevant regions in the solvated state. In our case, the restraining potential completely enveloped the binding pocket and was applied along the direction defined from the centers of mass of the whole protein and the binding pocket alone (aligned to the z-axis in the initial structure). We also applied a restraining bias to limit the translation of the center of mass of the protein (restrained a radius of 10 Å). We finally applied a restraining potential on the backbone of the residues that are not involved in the protein-ligand binding process (RMSD < 1 Å).

In our case, the MetaD simulations with these two LV-MetaD schemes in the WT MacroD2 systems led to very similar results, which further verified the reliability of both the shapes (Supplementary Table S1). Given that the ADPr binds to a buried and open active site, the MacroD1 and two MacroD2 variants systems were performed with parabolic-solid-shaped potentials. In the well-tempered MetaD simulation, Gaussian hills with a height of 1.2 kJ/mol was deposited every ps, and the Gaussian functions were rescaled using a bias factor of 20, while the widths of Gaussians were set as 1.0 Å, pi/8 radian, and 0.04 for the three CVs − *ρ*, θ, and σ, respectively. For all the systems, 0.8 μs of MetaD simulation were conducted and their convergence were achieved approximately at 0.6 μs (Supplementary Figure S3a). All the errors for free energy differences were evaluated using a block average analysis.

The lowest energy basin of each system has been equilibrated by unbiased MD. Flexibility of the binding site and the ligand atoms was evaluated and reported in Figure S11.

### Free energy calculation

The experimental absolute free energy values were calculated from the equilibrium dissociation constant Kd through the formula [36]

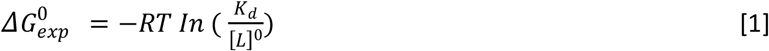

Where [L]^0^ is the standard state concentration of 1 mol/L, K_d_ is the ratio between the concentration in the bound state and that in the unbound state at specific temperature. In the MetaD simulation, the binding free energy (ΔG_metad) was the free-energy difference between the bound and unbound states.

However, the application of restraining potential caused a loss of the translational degrees of freedom of ligand in the solvated state, which would need to be corrected when calculating the absolute free energy (ΔG_standard) [2, 37, 38, 68].

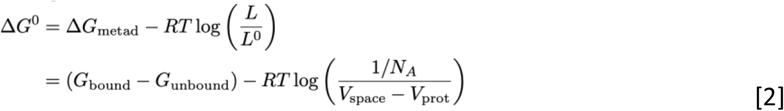

R is the gas constant, T the system temperature, [L] is the concentration of the ligand in the restraining space, N_A_ = 6.022×10^23^ is the Avogadro constant, V_space_ is the volume of the restraining potential, and V_prot_ is the volume of protein inside the restraint. The bound state was identified as the free energy minimum of the landscape and the unbound state as the area of the free energy surface with a distance between the protein center of mass and the ligand larger than 30 Å. Further information on the calculation of volume of parabolic-solid can be found in Figure S1.

## References

1. R Laurie, A.T., R.M.J.C.P. Jackson, and P. Science, Methods for the prediction of protein-ligand binding sites for structure-based drug design and virtual ligand screening. 2006. 7(5): p. 395–406.

2. Brotzakis, Z.F., V. Limongelli, M.J.J.o.C.T. Parrinello, and Computation, Accelerating the calculation of protein–ligand binding free energy and residence times using dynamically optimized collective variables. 2018. 15(1): p. 743–750.

3. Hu, G., H. Li, S. Xu, and J.J.I.j.o.m.s. Wang, Ligand Binding Mechanism and Its Relationship with Conformational Changes in Adenine Riboswitch. 2020. 21(6): p. 1926.

4. Vellore, N.A. and R.J.B.b. Baron, Molecular dynamics simulations indicate an induced-fit mechanism for LSD1/CoREST-H3-histone molecular recognition. 2013. 6(1): p. 1–9.

5. Chodera, J.D. and D.L. Mobley, Entropy-enthalpy compensation: role and ramifications in biomolecular ligand recognition and design. 2013.

6. Agafonov, R.V., C. Wilson, R. Otten, V. Buosi, D.J.N.s. Kern, and m. biology, Energetic dissection of Gleevec’s selectivity toward human tyrosine kinases. 2014. 21(10): p. 848–853.

7. Agafonov, R.V., C. Wilson, and D.J.F.i.m.b. Kern, Evolution and intelligent design in drug development. 2015. 2: p. 27.

8. Bernetti, M., M. Masetti, W. Rocchia, and A.J.A.r.o.p.c. Cavalli, Kinetics of drug binding and residence time. 2019. 70: p. 143–171.

9. Schuetz, D.A., W.E.A. de Witte, Y.C. Wong, B. Knasmueller, L. Richter, D.B. Kokh, et al., Kinetics for Drug Discovery: an industry-driven effort to target drug residence time. 2017. 22(6): p. 896–911.

10. Gobbo, D., V. Piretti, R.M.C. Di Martino, S.K. Tripathi, B. Giabbai, P. Storici, et al., Investigating Drug–Target Residence Time in Kinases through Enhanced Sampling Simulations. 2019. 15(8): p. 4646–4659.

11. Lee, K.S.S., J. Yang, J. Niu, C.J. Ng, K.M. Wagner, H. Dong, et al., Drug-target residence time affects in vivo target occupancy through multiple pathways. 2019. 5(9): p. 1614–1624.

12. Copeland, R.A.J.F.m.c., Conformational adaptation in drug–target interactions and residence time. 2011. 3(12): p. 1491–1501.

13. Miller, E., R. Murphy, D. Sindhikara, K. Borrelli, M. Grisewood, F. Ranalli, et al., A Reliable and Accurate Solution to the Induced Fit Docking Problem for Protein-Ligand Binding. 2020.

14. Gorgulla, C., A. Boeszoermenyi, Z.-F. Wang, P.D. Fischer, P.W. Coote, K.M.P. Das, et al., An open-source drug discovery platform enables ultra-large virtual screens. 2020. 580(7805): p. 663–668.

15. Iglesias, J., S. Saen-oon, R. Soliva, and V.J.W.I.R.C.M.S. Guallar, Computational structure-based drug design: Predicting target flexibility. 2018. 8(5): p. e1367.

16. Kokh, D.B., R.C. Wade, and W.J.W.I.R.C.M.S. Wenzel, Receptor flexibility in small-molecule docking calculations. 2011. 1(2): p. 298–314.

17. A Sotriffer, C.J.C.t.i.m.c., Accounting for induced-fit effects in docking: what is possible and what is not? 2011. 11(2): p. 179–191.

18. Guedes, I.A., F.S. Pereira, and L.E.J.F.i.p. Dardenne, Empirical scoring functions for structure-based virtual screening: applications, critical aspects, and challenges. 2018. 9: p. 1089.

19. Shan, Y., E.T. Kim, M.P. Eastwood, R.O. Dror, M.A. Seeliger, and D.E.J.J.o.t.A.C.S. Shaw, How does a drug molecule find its target binding site? 2011. 133(24): p. 9181–9183.

20. Pan, A.C., H. Xu, T. Palpant, D.E.J.J.o.c.t. Shaw, and computation, Quantitative characterization of the binding and unbinding of millimolar drug fragments with molecular dynamics simulations. 2017. 13(7): p. 3372–3377.

21. Wang, L., J. Chambers, and R. Abel, Protein–ligand binding free energy calculations with FEP+, in Biomolecular Simulations. 2019, Springer. p. 201–232.

22. Abel, R., L. Wang, E.D. Harder, B. Berne, and R.A.J.A.o.c.r. Friesner, Advancing drug discovery through enhanced free energy calculations. 2017. 50(7): p. 1625–1632.

23. de Oliveira, C., H.S. Yu, W. Chen, R. Abel, L.J.J.o.c.t. Wang, and computation, Rigorous free energy perturbation approach to estimating relative binding affinities between ligands with multiple protonation and tautomeric states. 2018. 15(1): p. 424–435.

24. Gapsys, V., S. Michielssens, D. Seeliger, and B.L.J.J.o.c.c. de Groot, pmx: Automated protein structure and topology generation for alchemical perturbations. 2015. 36(5): p. 348–354.

25. Samsudin, F., J.L. Parker, M.S. Sansom, S. Newstead, and P.W.J.C.c.b. Fowler, Accurate prediction of ligand affinities for a proton-dependent oligopeptide transporter. 2016. 23(2): p. 299–309.

26. Leonis, G., T. Steinbrecher, M.G.J.J.o.c.i. Papadopoulos, and modeling, A contribution to the drug resistance mechanism of Darunavir, Amprenavir, Indinavir, and Saquinavir complexes with HIV-1 protease due to flap mutation I50V: A systematic MM–PBSA and thermodynamic integration study. 2013. 53(8): p. 2141–2153.

27. Michielssens, S., J.H. Peters, D. Ban, S. Pratihar, D. Seeliger, M. Sharma, et al., A designed conformational shift to control protein binding specificity. 2014. 126(39): p. 10535–10539.

28. Gapsys, V., S. Michielssens, D. Seeliger, and B.L.J.A.C.I.E. de Groot, Accurate and rigorous prediction of the changes in protein free energies in a large-scale mutation scan. 2016. 55(26): p. 7364–7368.

29. Cournia, Z., B. Allen, W.J.J.o.c.i. Sherman, and modeling, Relative binding free energy calculations in drug discovery: recent advances and practical considerations. 2017. 57(12): p. 2911–2937.

30. Mobley, D.L. and M.K.J.A.r.o.b. Gilson, Predicting binding free energies: frontiers and benchmarks. 2017. 46: p. 531–558.

31. Leone, V., F. Marinelli, P. Carloni, and M.J.C.o.i.s.b. Parrinello, Targeting biomolecular flexibility with metadynamics. 2010. 20(2): p. 148–154.

32. Barducci, A., M. Bonomi, and M.J.W.I.R.C.M.S. Parrinello, Metadynamics. 2011. 1(5): p. 826–843.

33. Pfaendtner, J., Metadynamics to Enhance Sampling in Biomolecular Simulations, in Biomolecular Simulations. 2019, Springer. p. 179–200.

34. Capelli, R., A. Bochicchio, G. Piccini, R. Casasnovas, P. Carloni, M.J.J.o.c.t. Parrinello, et al., Chasing the full free energy landscape of neuroreceptor/ligand unbinding by metadynamics simulations. 2019. 15(5): p. 3354–3361.

35. Casasnovas, R., V. Limongelli, P. Tiwary, P. Carloni, and M.J.J.o.t.A.C.S. Parrinello, Unbinding kinetics of a p38 MAP kinase type II inhibitor from metadynamics simulations. 2017. 139(13): p. 4780–4788.

36. Bochicchio, A., G. Rossetti, O. Tabarrini, S. Krauβ, P.J.J.o.c.t. Carloni, and computation, Molecular view of ligands specificity for CAG repeats in anti-Huntington therapy. 2015. 11(10): p. 4911–4922.

37. Capelli, R., P. Carloni, and M.J.T.j.o.p.c.l. Parrinello, Exhaustive search of ligand binding pathways via volume-based metadynamics. 2019. 10(12): p. 3495–3499.

38. Limongelli, V., M. Bonomi, and M.J.P.o.t.N.A.o.S. Parrinello, Funnel metadynamics as accurate binding free-energy method. 2013. 110(16): p. 6358–6363.

39. Ibrahim, P. and T.J.C.O.i.S.B. Clark, Metadynamics simulations of ligand binding to GPCRs. 2019. 55: p. 129–137.

40. Cavalli, A., A. Spitaleri, G. Saladino, and F.L.J.A.o.c.r. Gervasio, Investigating drug–target association and dissociation mechanisms using metadynamics-based algorithms. 2015. 48(2): p. 277–285.

41. Saleh, N., P. Ibrahim, G. Saladino, F.L. Gervasio, T.J.J.o.c.i. Clark, and modeling, An efficient metadynamics-based protocol to model the binding affinity and the transition state ensemble of G-protein-coupled receptor ligands. 2017. 57(5): p. 1210–1217.

42. Lüscher, B., M. Bütepage, L. Eckei, S. Krieg, P. Verheugd, and B.H.J.C.r. Shilton, ADP-ribosylation, a multifaceted posttranslational modification involved in the control of cell physiology in health and disease. 2018. 118(3): p. 1092–1136.

43. Weixler, L., K. Schäringer, J. Momoh, B. Lüscher, K.L. Feijs, and R.J.N.A.R. Žaja, ADP-ribosylation of RNA and DNA: from in vitro characterization to in vivo function. 2021. 49(7): p. 3634–3650.

44. Haikarainen, T., M.M. Maksimainen, E. Obaji, and L.J.S.D.A.L.S.R. Lehtiö, Development of an inhibitor screening assay for mono-ADP-ribosyl hydrolyzing macrodomains using AlphaScreen technology. 2018. 23(3): p. 255–263.

45. Zhang, Y., M. Jumppanen, M.M. Maksimainen, S. Auno, Z. Awol, L. Ghemtio, et al., Adenosine analogs bearing phosphate isosteres as human MDO1 ligands. 2018. 26(8): p. 1588–1597.

46. Žaja, R., G. Aydin, B. Lippok, R. Feederle, B. Lüscher, and K.J.S.r. Feijs, Comparative analysis of MACROD1, MACROD2 and TARG1 expression, localisation and interactome. 2020. 10(1): p. 1–16.

47. Yang, X., Y. Ma, Y. Li, Y. Dong, L.Y. Lily, H. Wang, et al., Molecular basis for the MacroD1-mediated hydrolysis of ADP-ribosylation. 2020: p. 102899.

48. Chen, D., M. Vollmar, M.N. Rossi, C. Phillips, R. Kraehenbuehl, D. Slade, et al., Identification of macrodomain proteins as novel O-acetyl-ADP-ribose deacetylases. 2011. 286(15): p. 13261–13271.

49. Yang, X., Y. Ma, Y. Li, Y. Dong, L.Y. Lily, H. Wang, et al., Molecular basis for the MacroD1-mediated hydrolysis of ADP-ribosylation. 2020. 94: p. 102899.

50. Jankevicius, G., M. Hassler, B. Golia, V. Rybin, M. Zacharias, G. Timinszky, et al., A family of macrodomain proteins reverses cellular mono-ADP-ribosylation. 2013. 20(4): p. 508.

51. Neuvonen, M. and T.J.J.o.m.b. Ahola, Differential activities of cellular and viral macro domain proteins in binding of ADP-ribose metabolites. 2009. 385(1): p. 212–225.

52. Wazir, S., M.M. Maksimainen, and L.J.A.C.S.F.S.B.C. Lehtiö, Multiple crystal forms of human MacroD2. 2020. 76(10).

53. Invernizzi, M. and M.J.T.j.o.p.c.l. Parrinello, Rethinking metadynamics: from bias potentials to probability distributions. 2020. 11(7): p. 2731–2736.

54. Bonomi, M., G. Bussi, C. Camilloni, G.A. Tribello, P. Banáš, A. Barducci, et al., Promoting transparency and reproducibility in enhanced molecular simulations. 2019. 16(8): p. 670–673.

55. Pettersen, E.F., T.D. Goddard, C.C. Huang, G.S. Couch, D.M. Greenblatt, E.C. Meng, et al., UCSF Chimera—a visualization system for exploratory research and analysis. 2004. 25(13): p. 1605–1612.

56. Sastry, G.M., M. Adzhigirey, T. Day, R. Annabhimoju, and W.J.J.o.c.-a.m.d. Sherman, Protein and ligand preparation: parameters, protocols, and influence on virtual screening enrichments. 2013. 27(3): p. 221–234.

57. Lindorff-Larsen, K., S. Piana, K. Palmo, P. Maragakis, J.L. Klepeis, R.O. Dror, et al., Improved side-chain torsion potentials for the Amber ff99SB protein force field. 2010. 78(8): p. 1950–1958.

58. Jorgensen, W.L., J. Chandrasekhar, J.D. Madura, R.W. Impey, and M.L.J.T.J.o.c.p. Klein, Comparison of simple potential functions for simulating liquid water. 1983. 79(2): p. 926–935.

59. Wang, J., R.M. Wolf, J.W. Caldwell, P.A. Kollman, and D.A.J.J.o.c.c. Case, Development and testing of a general amber force field. 2004. 25(9): p. 1157–1174.

60. Jakalian, A., D.B. Jack, and C.I.J.J.o.c.c. Bayly, Fast, efficient generation of high-quality atomic charges. AM1-BCC model: II. Parameterization and validation. 2002. 23(16): p. 1623–1641.

61. Abraham, M.J., T. Murtola, R. Schulz, S. Páll, J.C. Smith, B. Hess, et al., GROMACS: High performance molecular simulations through multi-level parallelism from laptops to supercomputers. 2015. 1: p. 19–25.

62. Tribello, G.A., M. Bonomi, D. Branduardi, C. Camilloni, and G.J.C.P.C. Bussi, PLUMED 2: New feathers for an old bird. 2014. 185(2): p. 604–613.

63. Essmann, U., L. Perera, M.L. Berkowitz, T. Darden, H. Lee, and L.G.J.T.J.o.c.p. Pedersen, A smooth particle mesh Ewald method. 1995. 103(19): p. 8577–8593.

64. Hess, B., H. Bekker, H.J. Berendsen, and J.G.J.J.o.c.c. Fraaije, LINCS: a linear constraint solver for molecular simulations. 1997. 18(12): p. 1463–1472.

65. Bussi, G., D. Donadio, and M.J.T.J.o.c.p. Parrinello, Canonical sampling through velocity rescaling. 2007. 126(1): p. 014101.

66. Parrinello, M. and A.J.J.o.A.p. Rahman, Polymorphic transitions in single crystals: A new molecular dynamics method. 1981. 52(12): p. 7182–7190.

67. Moon, P. and D.E. Spencer, Field theory handbook: including coordinate systems, differential equations and their solutions. 2012: Springer.

68. Allen, T.W., O.S. Andersen, and B.J.P.o.t.N.A.o.S. Roux, Energetics of ion conduction through the gramicidin channel. 2004. 101(1): p. 117–122.

