## supplementary meterials for "An Enhanced Sampling Approach to the Induced Fit Docking Problem in Protein-Ligand Binding: the case of mono-ADP-ribosylation hydrolases inhibitors"

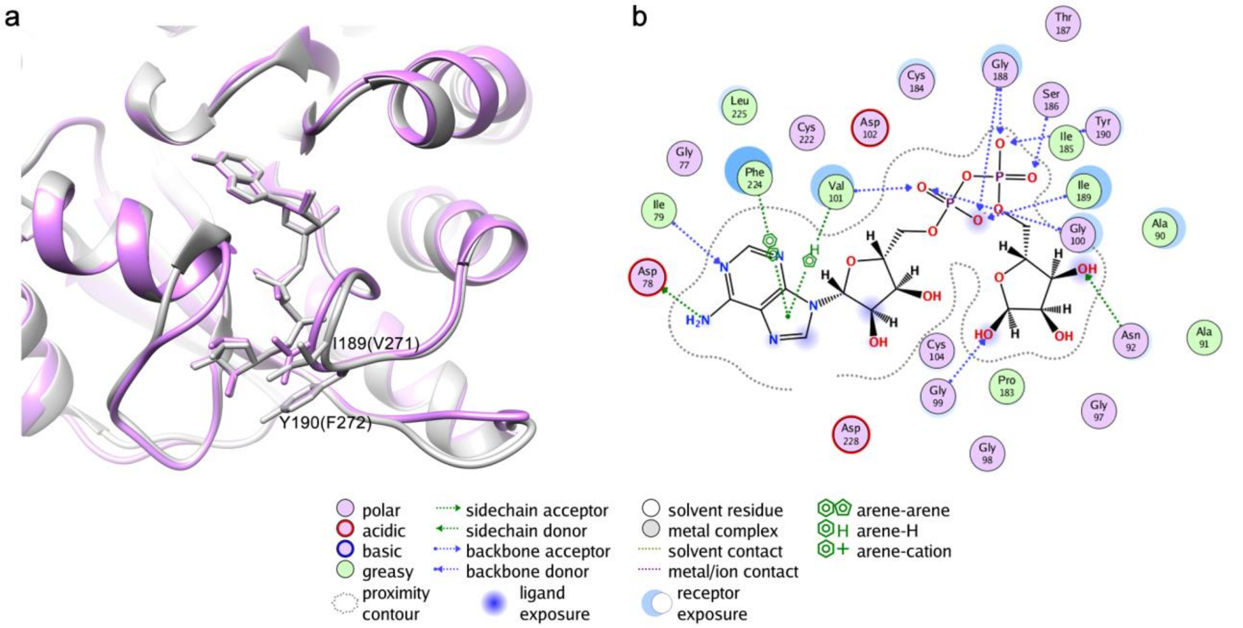


*Figure S1***.** *Superposition of holo-MacroD2 (in grey) and holo-MacroD1 crystal structure (in purple) (a) and the 2D plots of molecular interactions between MacroD2 and ligand ADPr in the holo-crystal structure (b).*

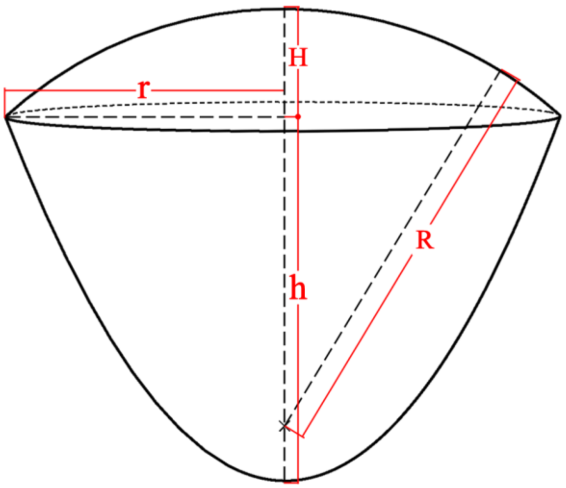


*Figure S2. Schematic diagram of parabolic-solids.*

The parabolic solid is comprised of the parabolic and spherical parts

$$V_{space} = V_{para} + V_{sphere} \tag{1}$$

among which the volume of the parabolic part can be calculated as

$$V_{para} = \int_{-\sigma^2/2}^{h-\sigma^2/2} \pi\sigma^2(2z + \sigma^2)dz \tag{2}$$

Where σ is the variables forming confocal paraboloids, h is the height of confocal paraboloids.

among which the volume of the spherical part can be calculated as

$$V_{sphere} = \int_{R-H}^{R} \pi(R^2 - z^2)dz \tag{3}$$

Where R is the restraint radius of sampling - the distance between the mass center of protein and ligand. H is the height of the spherical part.

*Table S1. Comparison of absolute free energy (ΔG _standard) for the WT MacroD2 and ADPr with two LV-MetaD schemes.*

| Scheme | $V_{prot}$ [a] (Å$^3$) | $V_{space}$ [b] (Å$^3$) | ΔG_correction (kcal/mol) | ΔG_metad (kcal/mol) | ΔG_standard (kcal/mol) |
|---|---|---|---|---|---|
| ⅓ sphere-solid | 11300 | 59864.8 | 2.0 | -7.3±0.3 | -9.3±0.3 |
| parabolic-solid | 14500 | 38222.7 | 1.6 | -7.8±0.3 | -9.4±0.3 |

*[a]$V_{prot}$, the volume of protein inside the restraining space, is calculated with VMD (probe radius = 0.5 Å, searching step = 0.5 Å, VDW = 1.7 Å). [b]$V_{space}$, the volume of the restraining potential, is obtained with mathematical calculation. The calculation of parabolic-solids is shown as follows.*

*Table S2. Distances between the center-of-mass of loop1 and loop2 in the crystal structures and in the binding process of four bias systems.*

| System | *holo*-structure (Å) | *apo*-structure (Å) | Maximum (Å) | Minimum (Å) | Bound State (Å) |
|---|---|---|---|---|---|
| MacroD2_para | 11.6 | 12.9 | 14.5 | 10.0 | 10.8 (basin 2) |
| MacroD1_para | 10.9 | 14.6 | 16.8 | 10.3 | 10.5 (basin 1) |
| MacroD2(Tyr190Asn) | / | / | 22.5 | 10.2 | 14.8 (basin 1) |
| MacroD2(Ile189Arg) | / | / | 17.5 | 9.8 | 10.6 (basin 1) |

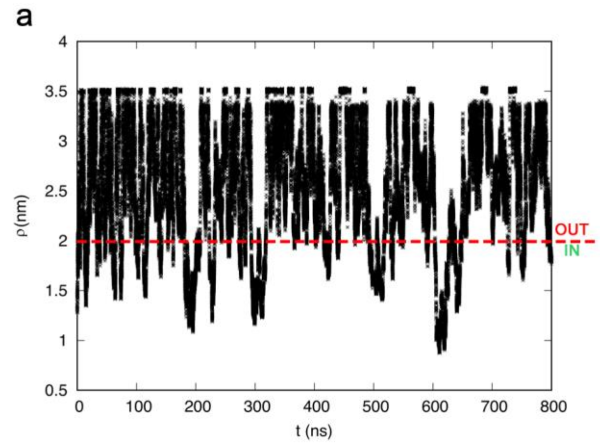


*Figure S3**.** Plot of the distance of center of mass between MacroD2 and ADPr along simulation (a) with parabolic-solid LV-MetaD scheme, (b) with ⅓ sphere-solid LV-MetaD scheme. Several recrossing events between bound (IN) and unbound (OUT) states can be observed.*

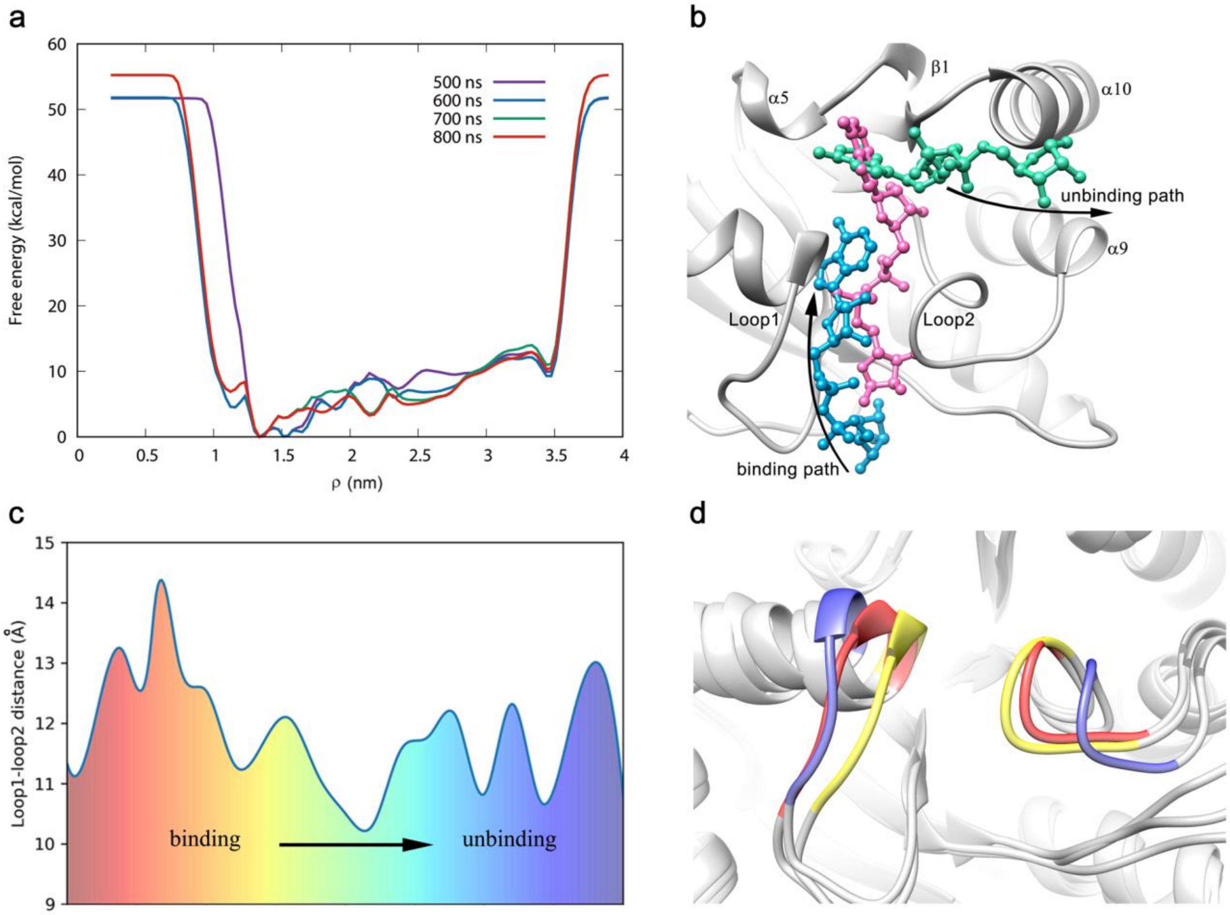


*Figure S4**.** The bias simulation of ligand ADPr binding to the WT holo-MacroD2 with parabolic-solid LV-MetaD scheme. (a) Binding free energy profiles of ADPr reconstructed along the ρ. After 600 ns the bound and unbound states were already sampled by the metadynamics. (b)The binding/unbinding pathway corresponding to the binding pose of basin 2. We show 3 representative ligand conformations: the first one (blue) is relative to the ligand when it is moving toward the binding pocket. The second one (pink) is the equilibrium conformation, and the third one (green) is relative to the step of the unbinding process. (c) Distribution of the distance between the center-of-mass of loop1 and loop2 in the binding/unbinding process of basin 2. (d) Movements of loop1 and loop2 in the binding/unbinding process of basin 2. Loop1 and loop2 in the holo-MacroD2 crystal structure is depicted in the red, narrowest loops during the process is in the yellow and the widest one is in the blue.*

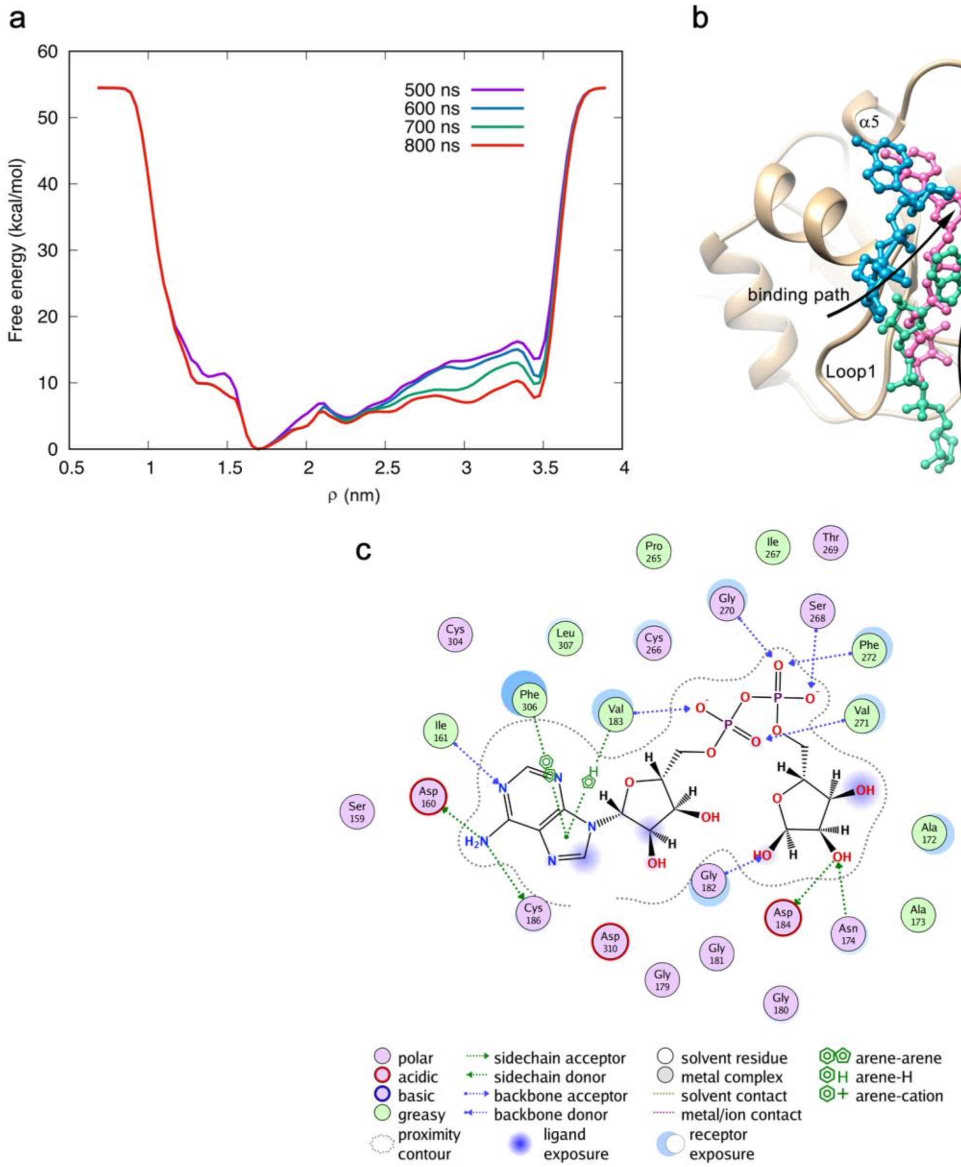


*Figure S5. The bias simulation of ligand ADPr binding to the apo-MacroD1 with parabolic-solid LV-MetaD scheme. (a) Binding free energy profiles of ADPr reconstructed along the ρ. After 500 ns the bound and unbound states were already sampled by the metadynamics. (b) The binding/unbinding pathway corresponding to the binding pose of basin 1. We show 3 representative ligand conformations: the first one (blue) is relative to the ligand when it is moving toward the binding pocket. The second one (pink) is the equilibrium conformation of basin 1, and the third one (green) is relative to the step of the unbinding process. (c) The 2D plots of molecular interactions between MacroD1 and ligand ADPr in the holo-crystal structure.*

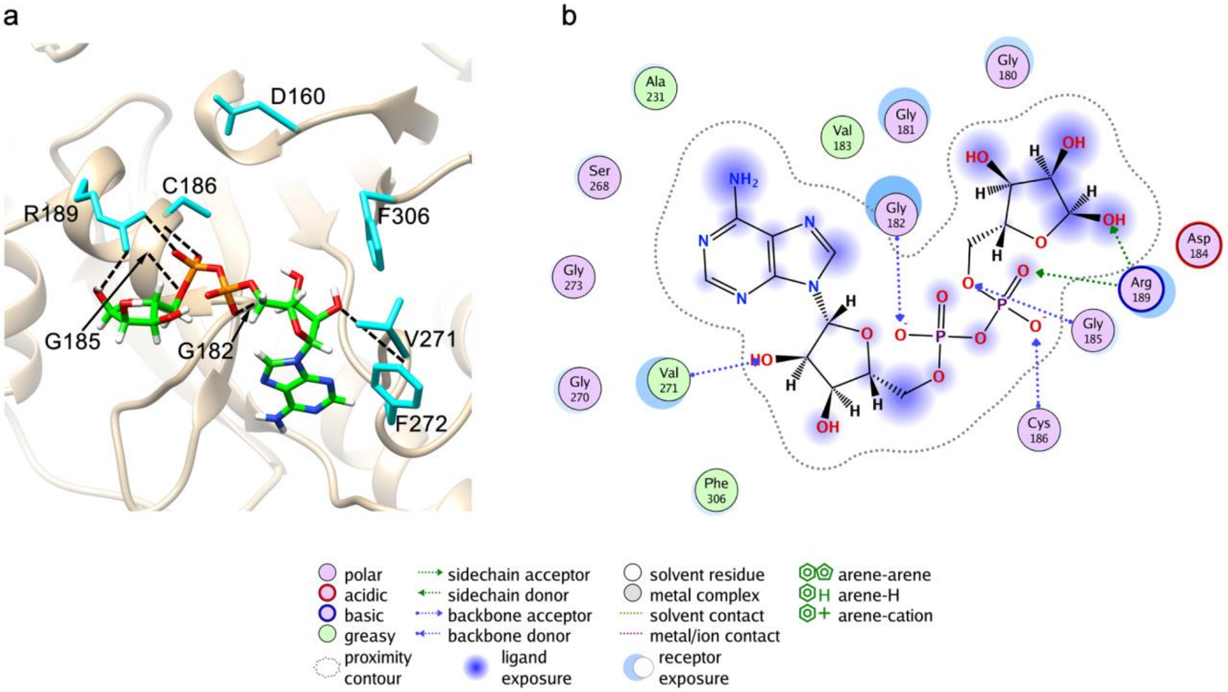


*Figure S6. (a) The representative conformation of free energy basin 2 in the bias simulation of ligand ADPr binding to the apo-MacroD1 and (b) its ligand-protein interaction diagram.*

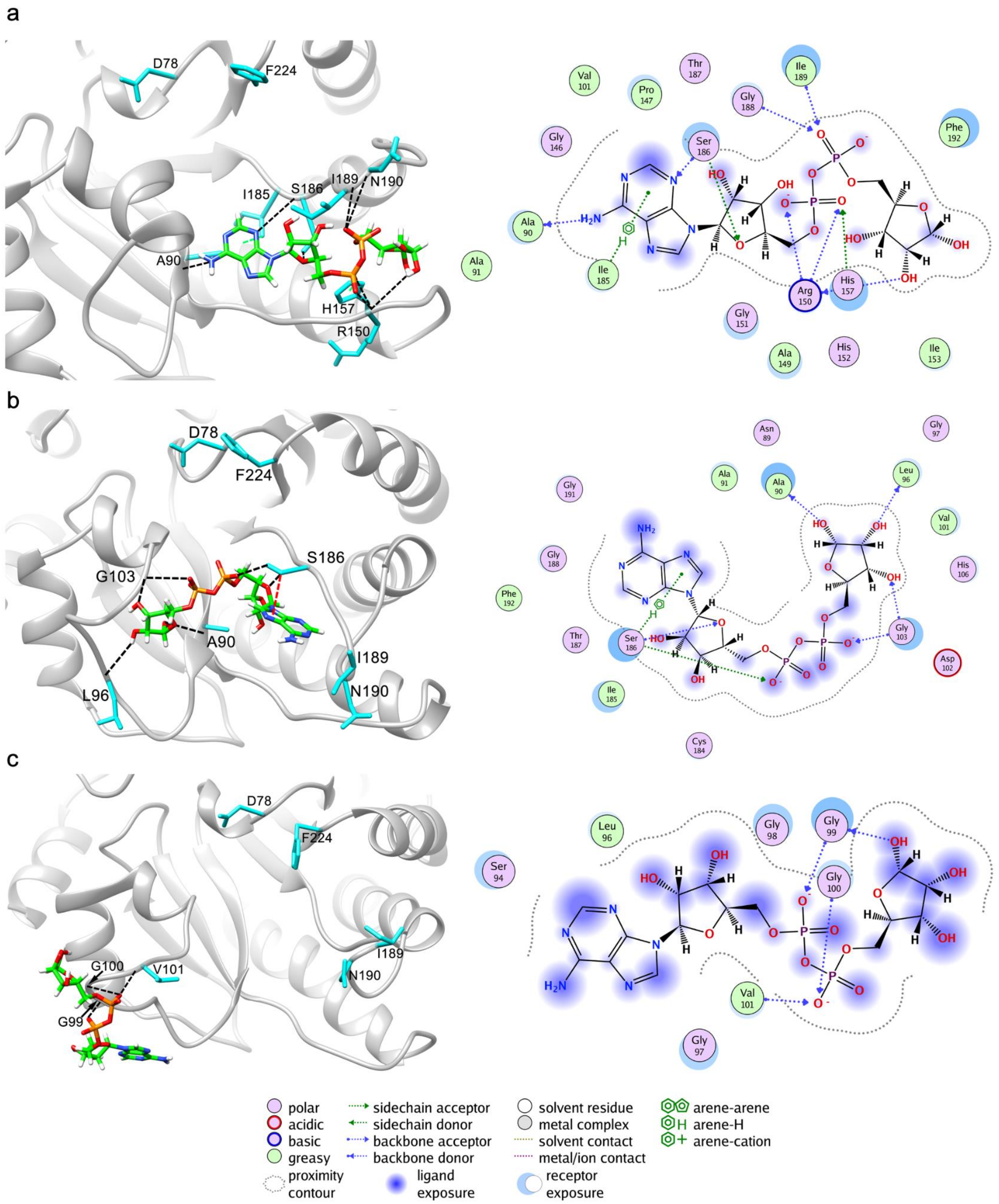


*Figure S7. The representative conformations and interaction diagrams of free energy basins in the Y190N MacroD2 system. (a) free energy basin 2. Bulk water is not shown for clarity purposes. Hydrogen bonds and hydrophobic interactions are depicted as black and green dashed lines, respectively. (b) free energy basin 3. (c) free energy basin 4.*

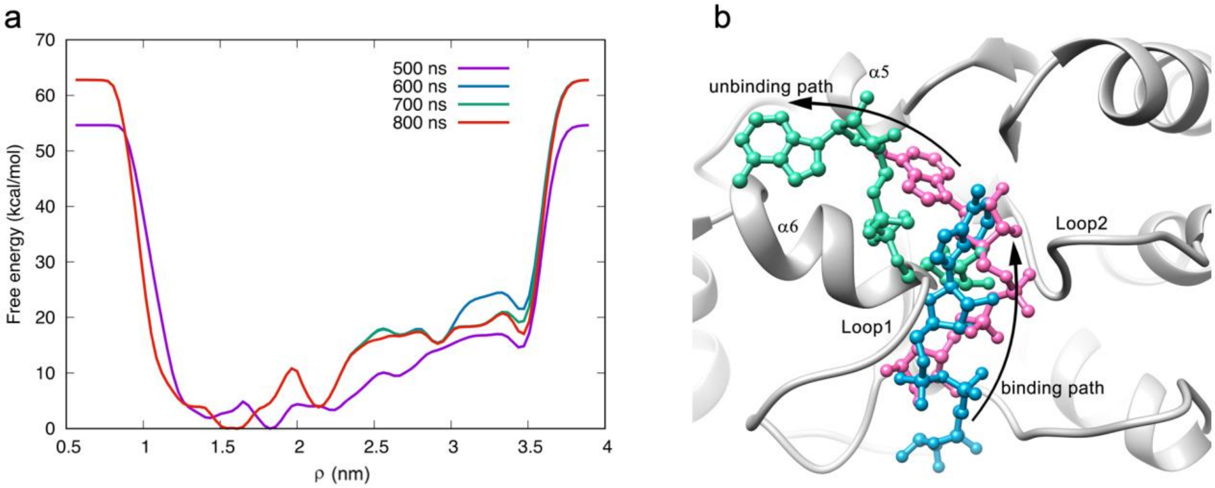


*Figure S8. The bias simulation of ligand ADPr binding to the mutant Y190N MacroD2 with parabolic-solid LV-MetaD scheme. (a) Binding free energy profiles of ADPr reconstructed along the ρ. After 500 ns the bound and unbound states were already sampled by the metadynamics. (b) The binding/unbinding pathway corresponding to the binding pose of basin 1. We show 3 representative ligand conformations: the first one (blue) is relative to the ligand when it is moving toward the binding pocket. The second one (pink) is the equilibrium conformation of basin 1, and the third one (green) is relative to the step of the unbinding process.*

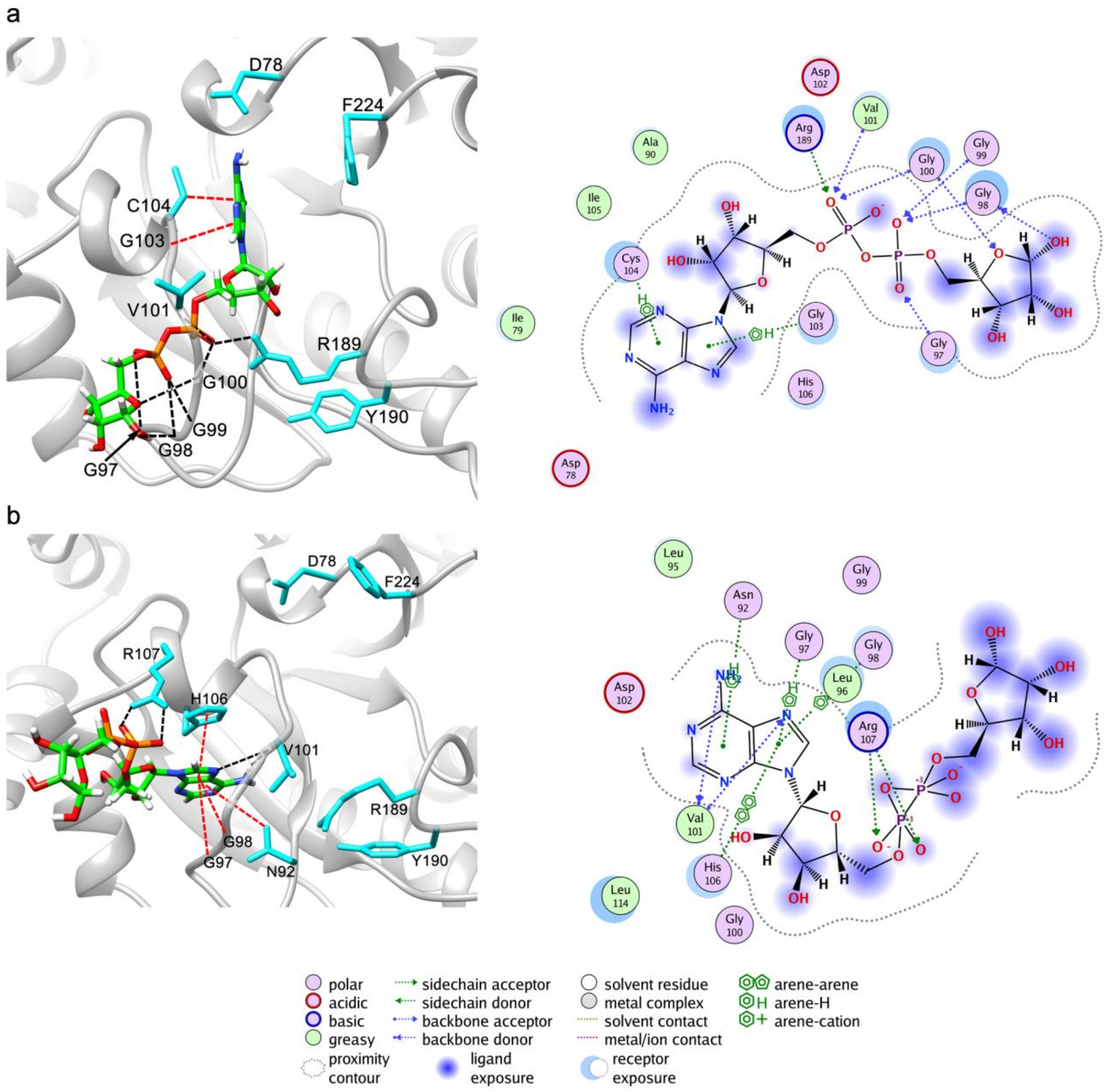


*Figure S9. The representative conformations and interaction diagrams of free energy basins in the I189R MacroD2 system. (a) free energy basin 2. Bulk water is not shown for clarity purposes Hydrogen bond and hydrophobic interactions are depicted as black and green dashed lines, respectively. (b) free energy basin 3.*

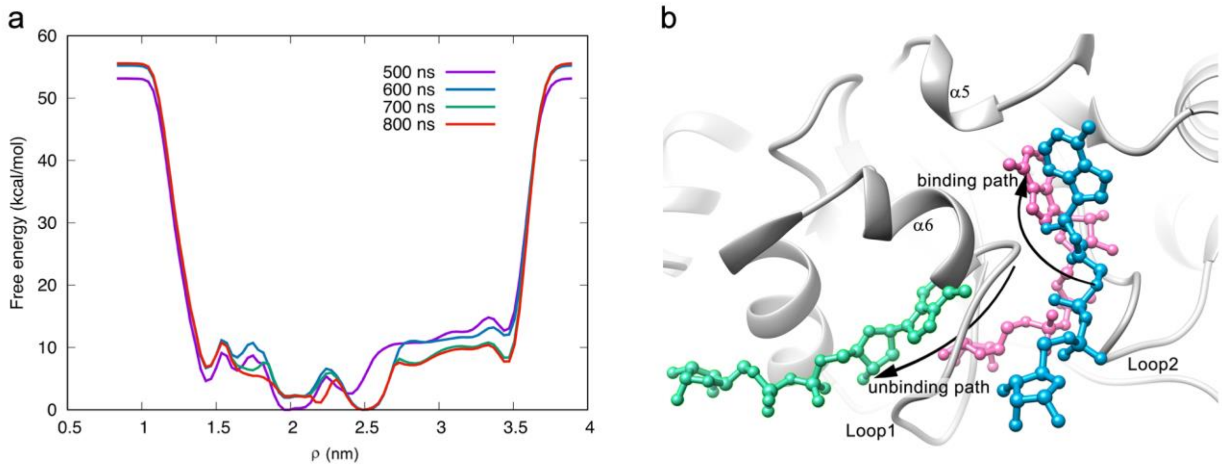


*Figure S10. The bias simulation of ligand ADPr binding to the mutant I189R MacroD2 with parabolic-solid LV-MetaD scheme. (a) Binding free energy profiles of ADPr reconstructed along the ρ. After 600 ns the bound and unbound states were already sampled by the metadynamics. (b) The binding/unbinding pathway corresponding to the binding pose of basin 1. We show 3 representative ligand conformations: the first one (blue) is relative to the ligand when it is moving toward the binding pocket. The second one (pink) is the equilibrium conformation of basin 1, and the third one (green) is relative to the step of the unbinding process.*

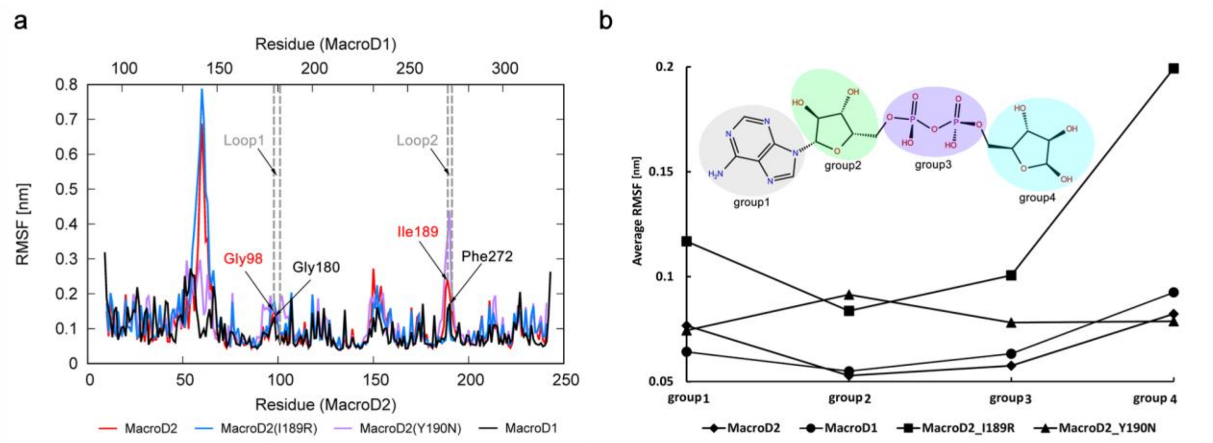


*Figure S11. Root mean square fluctuation (RMSF) of the structures of basin 1 through the equilibrium phase of MD simulation. (a) RMSF of protein backbone as function of residue number. The most flexible residues have been highlighted, red ones for the MacroD2 while black ones for the MacroD1. (b) Average RMSF of ADPr backbone as function of chemical groups.*
